# Receptor-enriched analysis of functional connectivity (REACT) for understanding cannabinoid neuropsychopharmacology

**DOI:** 10.1101/2025.06.02.657387

**Authors:** Ekaterina Shatalina, Natalie Ertl, Kat Petrilli, Shelan Ofori, Anna Borisova, Claire Mokrysz, H. Valerie Curran, Will Lawn, Tom P Freeman, Oliver Howes, Matthew B. Wall

## Abstract

**Background:** Cannabis is one of the most widely used psychoactive substances in the world and is increasingly investigated as a treatment for neuropsychiatric conditions. Cannabis contains delta-9-tetrahydrocannabinol (THC), which is thought to underlie its main psychoactive effects, and cannabidiol (CBD), which has been proposed to modulate the effects of THC on the brain. Both have activity at CB1 receptors, whilst THC also binds to CB2 receptors. However, it is unclear how THC’s effects on brain function are related to cannabinoid receptors, or how CBD co-administration affects this. We aimed to investigate the effects of vaporised THC and the moderating effects of CBD on CB1 and CB2 receptor-enriched functional connectivity.

**Methods:** Forty-eight participants (24 adolescents and 24 adults) who used cannabis 0.5–3 days/week (mean=1.5 days/week) participated in a randomised, crossover, placebo-controlled, double-blind experiment where they inhaled vaporised cannabis containing either THC-only (8mg/75kg person), THC+CBD (8 mg THC + 24 mg CBD/75 kg person) or no psychoactive compounds (placebo). Resting-state functional MRI data were collected approximately 50 minutes post-administration. Receptor distribution maps were derived from positron emission tomography (PET) imaging for CB1 receptors and the Allen Human Brain Atlas (AHBA) for CB2 receptors. Receptor-Enriched Analysis of Connectivity by Targets (REACT) analytical methodology was used to investigate changes in functional connectivity related to specific receptor targets.

**Results:** Inhalation of both THC-only and THC+CBD cannabis induced significant decreases in CB1 and CB2 receptor-enriched functional connectivity compared to placebo. The THC+CBD condition showed more extensive reductions than THC-only cannabis for both CB1- and CB2- enriched functional connectivity, affecting regions including the dorsolateral prefrontal cortex, cingulate cortex, insula, hippocampus, amygdala, and putamen. Higher THC plasma levels were associated with greater decreases in functional connectivity in the THC+CBD condition. Exploratory analyses identified significant positive and negative relationships between subjective drug effects and receptor-enriched functional connectivity in a region-dependent manner.

**Conclusions:** Cannabis with and without CBD decreased resting-state functional connectivity in networks associated with CB1 and CB2 receptor distribution. Co-administration of CBD with THC appears to enhance these effects. The REACT analytical methodology identified changes in CB1- and CB2-enriched functional connectivity are related to THC and CBD plasma levels as well as subjective drug effects across a range of regions.

## Introduction

The pharmacology of cannabis is complex due to the presence of multiple cannabinoids that can interact with each other and act on a range of targets. Of these, delta-9- tetrahydrocannabinol (THC) and cannabidiol (CBD) are two major, naturally occurring cannabinoids that may exhibit polypharmacological effects when administered together. THC produces most of the psychotropic effects of cannabis, mostly through partial agonism of the Cannabinoid receptor 1 (CB1) (*K*_i_ = 5.05–80.3 nM) (Pertwee 2008). CB1 is the most abundant G protein-coupled receptor in the brain, widely expressed in neurons throughout the central nervous system and is primarily localised to presynaptic terminals, where it regulates neurotransmitter release through inhibition of adenylyl cyclase and voltage-gated calcium channels (Mackie 2008). Activation of CB1 by THC underpins THC’s acute psychotropic effects: euphoria, appetite stimulation, memory impairment, and analgesia (Pertwee 2008).

THC also has activity at CB2 receptors (*K*_i_ = 1.73–75.3 nM), where it acts as a partial agonist (Pertwee 2008). Initially characterised as a receptor involved in peripheral immune responses, CB2 is now suggested to play a role in the central nervous system. CB2 is expressed at low levels in neurons and glial cells in the healthy brain, and its activation exerts anti-inflammatory effects by reducing the release of pro-inflammatory cytokines and modulating microglial activity (for review, see (Grabon, Rheims et al. 2023)).

Evidence also suggests CB2 activation modulates synaptic transmission *in vitro* (Kano, Ohno-Shosaku et al. 2009, Morgan, Stanford et al. 2009, Atwood, Straiker et al. 2012, Franklin and Carrasco 2012, Hwang, Kim et al. 2020, Sadanandan, Kreko-Pierce et al. 2020) and decreases neuronal excitability both *in vitro* (den Boon, Chameau et al. 2012, Zhang, Gao et al. 2014, Zhang, Gao et al. 2017, Ma, Gao et al. 2019, Yu, Liu et al. 2021, Zhang, Shen et al. 2021) and *in vivo* (Sokal, Elmes et al. 2003, Elmes, Jhaveri et al. 2004, Nackley, Zvonok et al. 2004, Jhaveri, Elmes et al. 2008, Zhang, Gao et al. 2014, Stempel, Stumpf et al. 2016). Rodent studies further suggest CB2R activation modulates feeding, motor function, memory, learning and mood (reviewed in (Grabon, Rheims et al. 2023)). The role of CB2 in mediating the psychotropic effects of THC remains poorly understood.

CBD, in contrast, acts as a negative allosteric modulator of CB1 receptors (Kb = 0.27-0.35µM, Ki>4µM) (Laprairie, Bagher et al. 2015), and there is evidence to show it has anxiolytic, antipsychotic, and anti-addictive properties (White 2019). CBD has been hypothesised to be oppositional to THC in its actions (Curran, Freeman et al. 2016, Wall, Pope et al. 2019), however, studies co-administering THC and CBD have produced mixed results dependent on factors including routes of administration, relative doses and timing of assessments (Freeman, Petrilli et al. 2019). The most recent evidence comes from a study that co-administered inhaled CBD and THC in different ratios ranging from 0:1 to 3:1 CBD:THC to healthy volunteers in a double-blind, within-subject, randomised manner (Englund, Oliver et al. 2023). The study reports that CBD did not modulate the effects of THC on cognitive and psychotic side effects, even at the highest dose, suggesting it does not have protective effects against the acute adverse effects of cannabis (Englund, Oliver et al. 2023).

Studies using fMRI following acute inhaled cannabinoid administration can provide insight into the neuropsychopharmacology of THC and CBD. For example, a network connectivity analysis of the present study, which included 24 adolescents and 24 adults, found that THC reduced functional connectivity compared to placebo (Ertl, Freeman et al. 2024). The authors found that coadministration of CBD with THC may enhance the effects of THC, resulting in a greater reduction in functional connectivity, in regions of the executive control and salience networks, than that caused by administration of THC alone (Ertl, Freeman et al. 2024). In contrast to this, a prior study using a similar design showed that CBD restored disruption of the salience network by THC in a group of 17 adults (Wall, Pope et al. 2019).

Informing drug-induced changes in brain function with biological information is an active area of research. For example, one study found that the COMT genotype modulated fMRI responses to THC, suggesting prefrontal dopamine levels may relate to the acute effects of the drug. (Pelgrim, Ramaekers et al. 2023)

However, a more powerful approach to this is Receptor-Enriched Analysis of Connectivity by Targets (REACT) (Dipasquale, Selvaggi et al. 2019). This method evaluates drug-induced changes in functional connectivity that relate to the receptor target(s) of the drug by integrating data on the spatial distribution of the drug target into the analyses. For example, by using a reference image produced by Positron Emission Tomography (PET) with a tracer that binds to a relevant target (Dipasquale, Selvaggi et al. 2019, Dipasquale, Martins et al. 2020, Lawn, Dipasquale et al. 2022). In summary, REACT uses maps of receptor expression, which allows researchers to explore the effects of drugs on functional connectivity in brain networks related to those receptors, deriving a measure of brain function related to the drug acting on a specific receptor target.

Previous work using the method has linked receptor-enriched connectivity to subjective drug effects and drug plasma levels. For example, MDMA-induced functional connectivity changes were found to be primarily mediated by the serotonin transporter (5-HTT) and 5-HT1A receptor, with MDMA plasma levels correlating with connectivity increases in 5-HT1A- enriched maps and MDMA-induced decreases in 5-HT2A-enriched connectivity correlating with subjective experience of spirituality (Dipasquale, Selvaggi et al. 2019). Similarly, work on LSD showed that while 5HT2a activity is central to changes in connectivity, LSD also alters 5HT1a, 5HT1b, D1, and D2 receptor-enriched functional connectivity, with serotonergic systems driving perceptual effects and dopaminergic systems modulating cognition and selfhood (Lawn, Dipasquale et al. 2022).

Human PET data availability is limited as only a restricted number of ligands are available, and therefore, most receptor targets in the brain cannot be imaged non-invasively, however, the Allen Human Brain Atlas (AHBA) is a unique resource that provides whole-brain genome-wide transcriptomic data suitable for REACT analyses (Shen, Overly et al. 2012). The atlas contains expression levels of >20,000 genes profiled by ∼60,000 microarray probes across different brain regions. Importantly, the data are spatially resolved in the standard Montreal Neurological Institute (MNI) coordinate system, allowing for a one-to-one spatial matching with neuroimaging data (Shen, Overly et al. 2012, Selvaggi, Rizzo et al. 2021).

In our present study, we build on the functional connectivity work on this cohort published by Ertl et al. (2024) to explore the effects of THC-only and THC+CBD-containing cannabis on the CB1 and CB2 receptor-enriched functional connectivity in groups of young adults and adolescents. We additionally include validation analyses of using AHBA data for REACT analyses by comparing the results of analyses that use a PET-derived and AHBA-derived version of CB1 distribution maps.

## Methods

The data presented here is from the CannTeen-Acute study. The full study protocol, which includes details on objectives, data collection methods, tasks, and power calculations, is available online (https://osf.io/z638r/). This study was not considered a clinical trial by the UK Medicines and Health Care Products Regulatory Agency, however, it was registered on clinicaltrials.gov to follow good practice on April 20, 2021, with the ID NCT04851392 (https://www.clinicaltrials.gov/study/NCT04851392). Additional evaluation of baseline symptomatology of subjects included in this study and the acute effects on the primary outcomes can be found in (Lawn, Trinci et al. 2023). The current study includes analyses of data that have been previously reported in (Ertl, Freeman et al. 2024).

### Ethics approval

Ethical approval was obtained from the University College London Research Ethics Committee, project ID 5929/005. The study was conducted in line with the Declaration of Helsinki, and all participants provided written informed consent to participate.

### Participants

Full demographic and drug-history information is shown in Table 1. Subjects were 48 semi-regular cannabis users who used cannabis 0.5 to 3 days/week over the past three months. The sample included 24 adults (26–29 years, mean = 27.8 years, 12 females) and 24 adolescents (16–17 years, mean = 17.2 years, 12 females) recruited using online advertisements and word of mouth from the greater London area. Adult users were excluded if they had used cannabis regularly before the age of 18. Volunteers were excluded if there was a history of or current psychosis or an immediate family member with a psychotic disorder, as well as if there were any other medical problems considered clinically significant for the study. Additional drug-related exclusion criteria were previous negative experiences with cannabis, alcohol use >5 times per week and use of any other illicit drug >twice per month. All subjects were in good physical health and were not receiving treatment for any mental health condition. Full inclusion/exclusion criteria are detailed in (Lawn, Trinci et al. 2023).

**Table 1:**
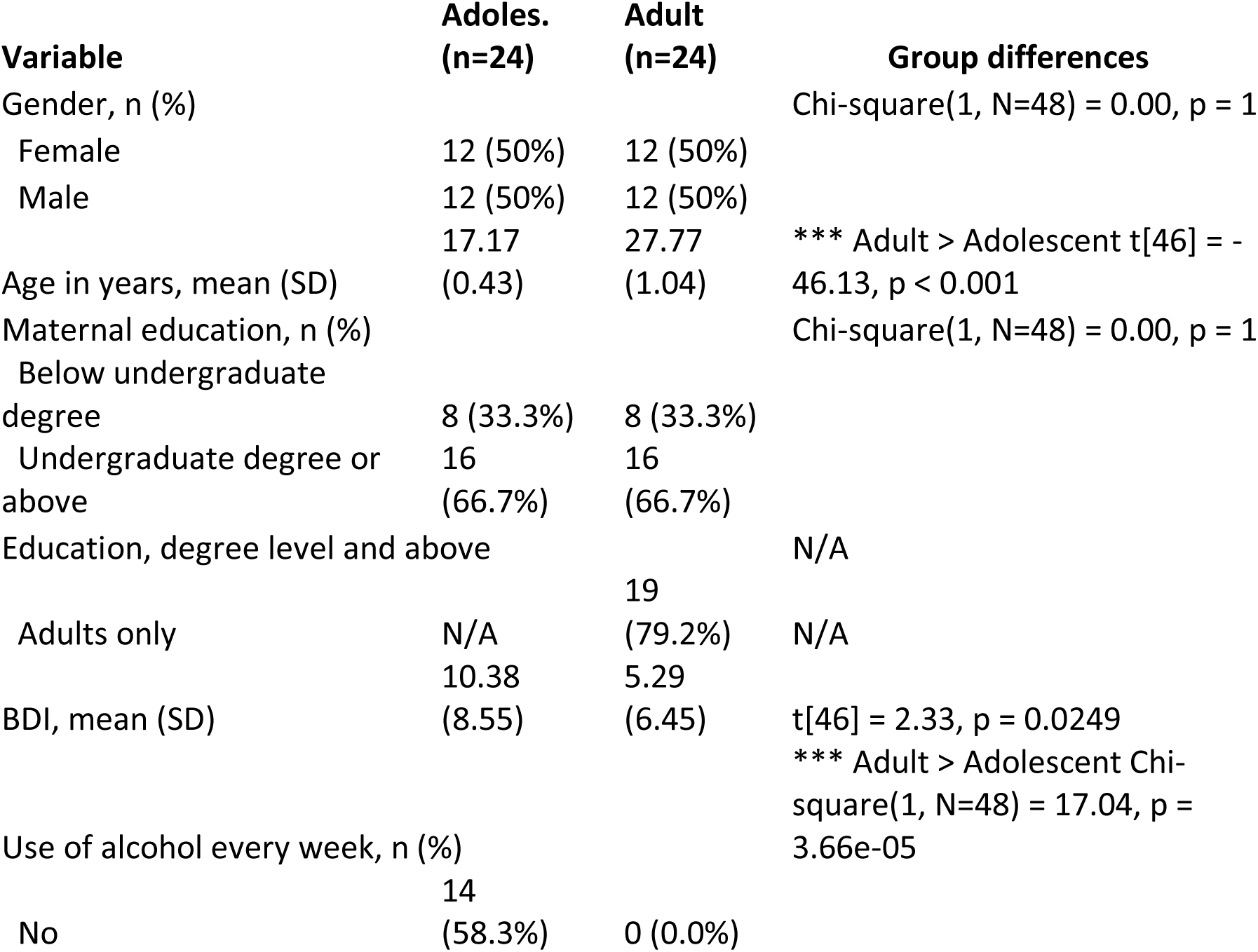
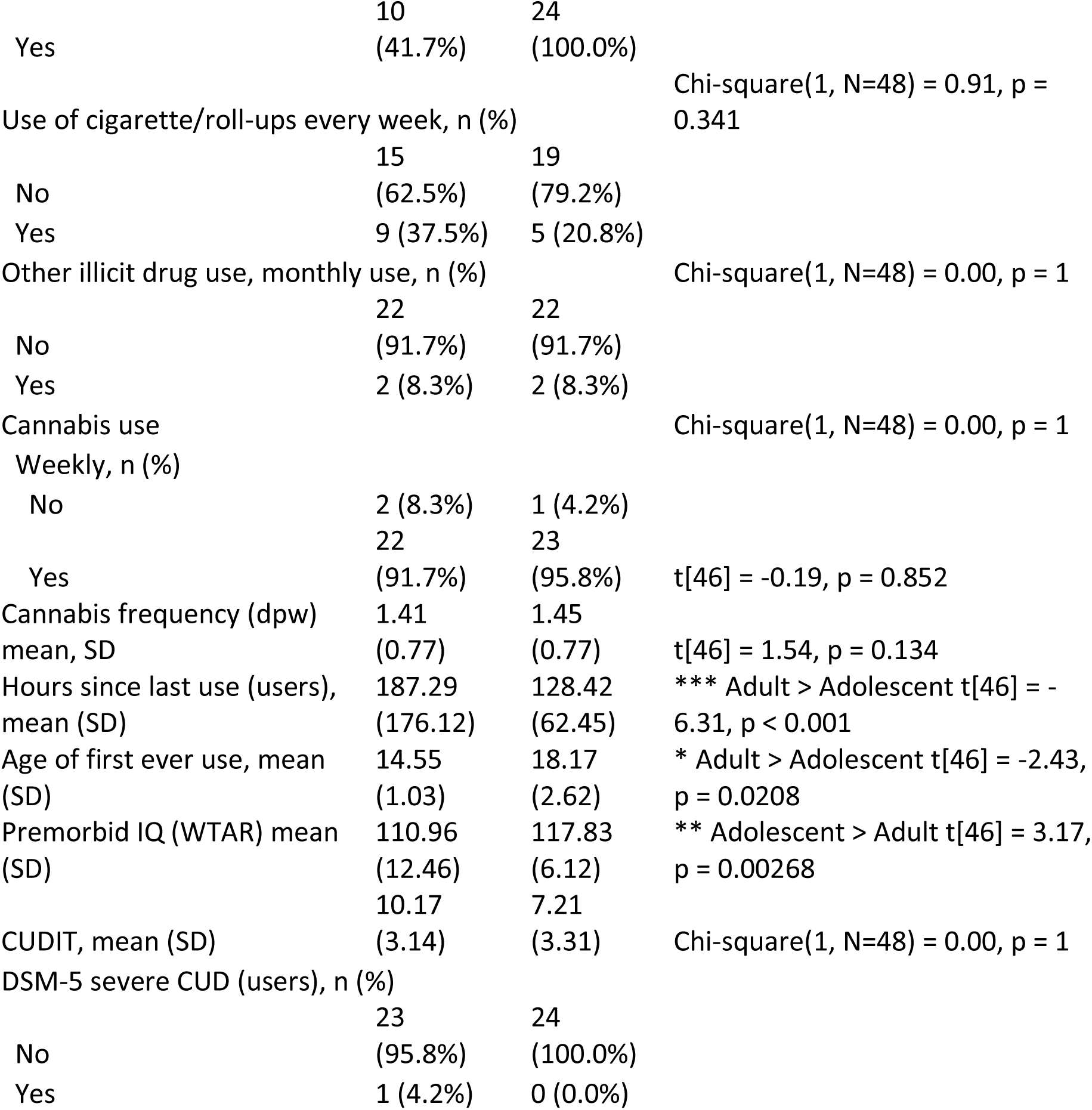
Sociodemographic characteristics of the full sample (n = 48). BDI is the Beck Depression Inventory. WTAR is Wechsler’s Test of Adult Reading. CUDIT is the Cannabis Use Disorder Inventory Test. Continuous data are presented as mean [SD], and categorical data are presented as n (%). Group differences are highlighted in the final column using appropriate tests for each data type (*χ*^2^ and t-tests; *P < 0.05, **P < 0.01, ***P < 0.001).

### Study design and scales

A randomised, crossover, placebo-controlled, double-blind design was used to compare cannabis containing THC but no CBD (THC-only) and cannabis containing THC and CBD (THC+CBD) with matched placebo cannabis containing neither compound. Participants were randomly assigned to one of three treatment order conditions based on a Latin Square design, in blocks of twelve participants. Drug order was balanced across participants and within age groups and gender.

Ratings of subjective effects using visual analogue scales (VAS) (scored from 0 to 10) were approximately 30 minutes after the drug administration period (after the MRI scan)(Lawn, Trinci et al. 2023). Behavioural variables of interest included the ‘Feel drug effect’ scale as it best captured total subjective drug effects, as well as scales for “paranoid”, “calm” and “anxious”.

### Drug administration

First, participants completed instant saliva drug (Alere DDSV 703 or ALLTEST DSD-867MET/C) and breathalyser tests (Lion Alcometer 500), as well as self-reported abstinence, to confirm no use of alcohol in the previous 24 hours and no use of cannabis or other illegal drugs in the previous 72 hours.

Cannabis (dried medical cannabis flower) was sourced from Bedrocan (Netherlands) and imported into the UK under a Schedule 1 Home Office License. The cannabis was administered with a Volcano Medic Vaporiser (Storz and Bickel) set at 210 °C. Three types of cannabis were used to create the formulations: Bedrocan (20.2% THC, 0.1% CBD), Bedrolite (0.4% THC, 8.5% CBD), and Bedrobinol (no THC or CBD). There was an absence of microbes, yeasts, aflatoxins, pesticides, and heavy metals in both Bedrolite and Bedrobinol, and there was the presence of cannabinol at 0.1%. Appropriate quantities of these three cannabis types were combined to produce the following treatments, matched for overall weight of cannabis: 0.107 mg/kg THC in the “THC only” condition (e.g. 8 mg THC for a 75 kg person), 0.107 mg/kg THC plus 0.320 mg/kg CBD in the “THC + CBD” condition (e.g. 8mg THC and 24 mg CBD for a 75 kg person), or placebo cannabis (0 mg THC, 0 mg CBD). The dose of THC used was equivalent to 1.6 standard units of THC (Freeman and Lorenzetti 2020). Subjects inhaled two balloons (each within nine minutes, a total of 18 min), with the experimenters monitoring standard timings for inhalation. This method of administration has been extensively used in previous work (Mokrysz, Freeman et al. 2016, Mokrysz, Shaban et al. 2021) and is safe and effective at delivering cannabinoids and producing well-documented behavioural and subjective effects. Placebo cannabis was closely matched to the active drug conditions in both appearance and smell, and all researchers present (as well as the participant) were blinded to the drug conditions. Additional staff (not present at the testing sessions) blinded the treatment conditions in advance of the testing sessions. The minimum washout period between testing sessions was three days, the mode was seven days, and the maximum was 51 days.

Participants received video training for the vaporiser during their baseline/screening session. Approximately 30 minutes following drug administration, participants were situated in the MRI scanner for a ∼one-hour scanning session. Blood samples were collected at -5, +20, +30, +160 minutes of cannabis administration as detailed in (Lawn, Trinci et al. 2023). For this sub-study, blood samples collected at +20 minutes were used as they reflected plasma concentrations closest to the resting state fMRI data acquisition. THC plasma levels were compared between THC-only and THC+CBD visits.

The resting-state scan was eight minutes long and was acquired towards the beginning of the scanning session, approximately 50 minutes after the start of drug administration. Previous work has shown that subjective effects of vaporised cannabis have a fast onset, and stay at a high level for approximately 60–90 min (Spindle, Cone et al. 2018). The timing of the resting-state scan was therefore close to the expected time of peak effects. Participants were instructed to keep their eyes open but blink as normal during the scan to mitigate against them falling asleep.

### MRI acquisition

Data were acquired on a Siemens Magnetom Verio 3T MRI scanner (n=38), with 10 subjects (5 teens, 5 adults) acquired on a Siemens Trio 3T MRI scanner (scanner type was included as a confound in later group-level analyses), both using 32-channel phased-array head coils. Subjects always completed all three scanning sessions on the same scanner to avoid additional intra-subject variability. Sequences used on the two scanners were harmonised to be identical, and these two particular scanners have been previously documented to produce highly similar results with EPI-BOLD sequences (Demetriou, Kowalczyk et al. 2018).

T_1_-weighted structural images were acquired using a Magnetisation Prepared Rapid Gradient Echo (MPRAGE) sequence (TR = 2300 ms, TE = 2.98 ms, flip angle = 9°, parallel imaging acceleration factor = 2), with a spatial resolution of 1 mm isotropic. T_2_* images were acquired using a multiband gradient echo Echo-Planar Imaging (EPI) sequence (TR = 1250 ms, echo time, TE = 30 ms, flip angle = 62°, multiband acceleration factor = 2, GRAPPA = 2, bandwidth = 1906Hz/pixel). This sequence was based on those previously documented and validated on both scanners by Demetriou et al. (2018). A total of 384 volumes were collected for each subject, with a field-of-view of 192 mm and a matrix size of 64 x 64 mm, yielding an in-plane resolution of 3 x 3 mm. Slice thickness was also 3 mm, resulting in isotropic voxels. Forty-four slices were collected using an interleaved acquisition. Phase encoding direction was anterior to posterior. The forebrain, midbrain, and hindbrain (including the cerebellum) were covered.

### Data preprocessing

Data preprocessing was carried out using FSL (FMRIB Software Library v6.0.0; http://www.fmrib.ox.ac.uk/fsl/) and included brain extraction of the anatomical data carried out using BET and the fsl_anat script was used for additional anatomical data pre-processing. Motion correction was performed with FMRIB Linear Image Registration Tool (MCFLIRT), with spatial smoothing using a Gaussian kernel of full width at half maximum (FWHM) 6mm. A two-step co-registration, first to the subject’s individual anatomical image, followed by registration to an anatomical template image in standard stereotactic space (MNI152), was performed, with temporal filtering applied at 100s. Time series from each subject’s white matter and cerebrospinal fluid were used as regressors of no interest alongside 24 head motion regressors. An extended evaluation of head motion in this dataset was carried out previously and is presented in the supplement of (Ertl, Freeman et al. 2024).

### Receptor-Enriched Analysis of Connectivity by Targets (REACT)

This analysis was focused on two receptor targets, CB1 and CB2 receptors. An atlas of CB1 receptor expression was obtained from the JuSpace toolbox (Dukart, Holiga et al. 2021). This image was derived from 10 averaged images acquired using [^11^C]OMAR positron emission tomography (Normandin, Zheng et al. 2015, Hansen, Shafiei et al. 2022). As no PET tracer is available for use in humans that binds to the CB2 receptor, we derived a CB2 receptor expression map from the Allen Human Brain Atlas (AHBA) downloaded from the Neurosynth website (neurosynth.org/genes/CNR2). These maps include normalised data on mRNA expression acquired post-mortem for six subjects across all cortical and subcortical regions displayed in MNI space. Previous work has shown strong concordance between PET and gene expression data obtained from the AHBA (Rizzo, Veronese et al. 2014, Hansen, Markello et al. 2022). However, to provide additional validation of using gene expression data for REACT-based analysis of fMRI data, we additionally derived the CB1 receptor expression map from the AHBA (neurosynth.org/genes/CNR1) to compare results to those using the PET-derived map.

The AHBA-derived receptor maps were processed to more closely resemble the CB1 PET atlas image we used in this study. Specifically, the gene expression images were smoothed with a 6mm FWHM kernel to match the smoothness of the PET data and normalised to have values ranging from 0–100 to match the CB1 PET-derived atlas map (Dukart, Holiga et al. 2021). Images were masked using a 75% grey matter mask derived from the MNI152 T1 structural template; resultant images are shown in Figure 1.

**Figure 1:**
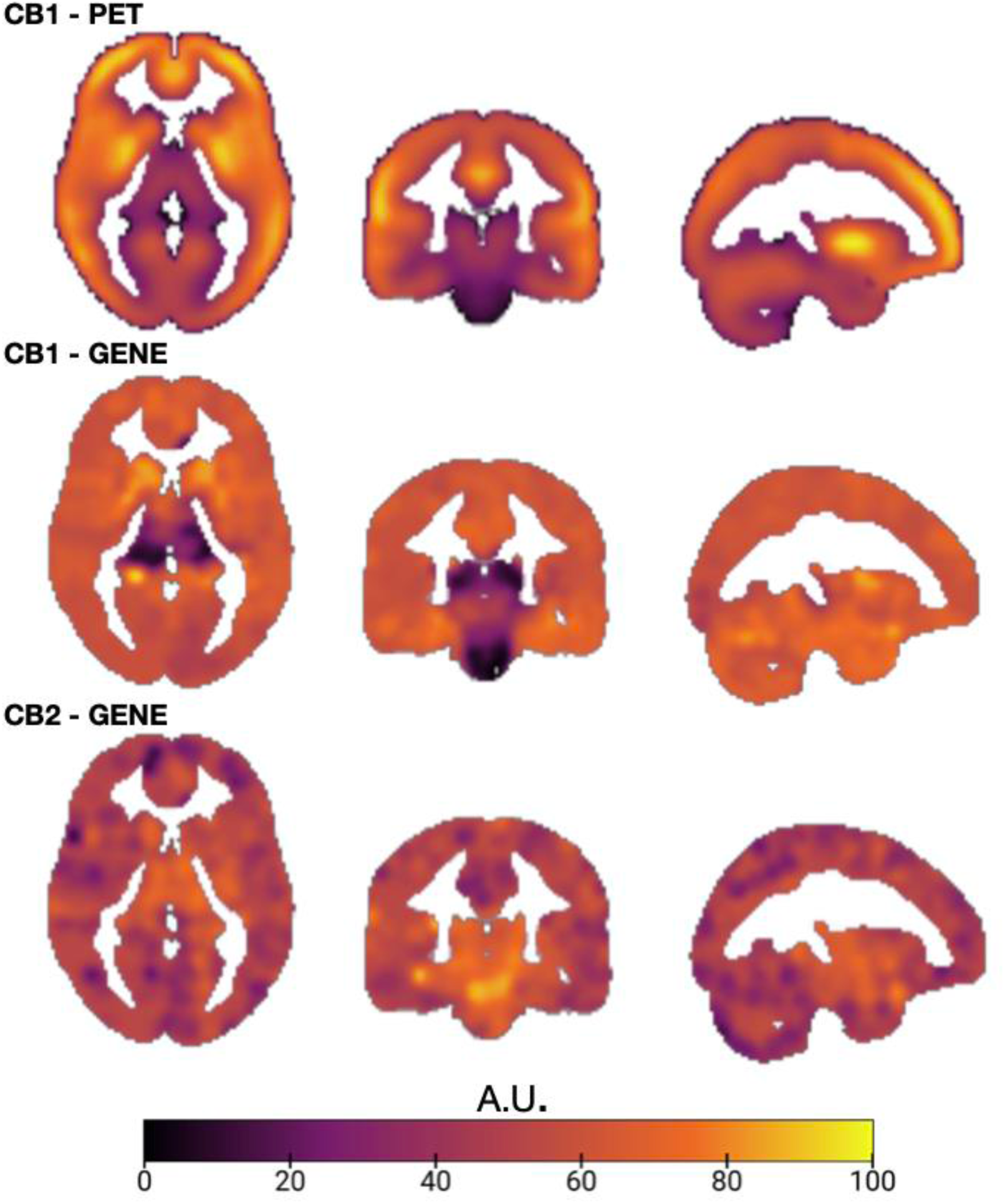
Spatial distribution of CB1 (derived from [^11^C]OMAR PET), CB1 (derived from the Allen Human Brain Atlas, AHBA) and CB2 (AHBA) expression maps displayed in MNI152 space with 75 % grey-matter masking applied, scaled 0-100. Warmer colours indicate higher relative receptor density.

REACT analyses closely followed work by Dipasquale et al., who first developed the analysis methodology (Dipasquale, Selvaggi et al. 2019). All target maps were entered into the first step of a dual regression analysis implemented using fsl_glm. This uses the receptor maps as a set of spatial regressors to estimate functional connectivity with voxels in the receptor maps, weighed by the level of target expression. The rs-fMRI data and the design matrix were demeaned to improve the model fit, as is standard for dual regression analyses (Filippini, MacIntosh et al. 2009, Smith, Utevsky et al. 2014, Nickerson, Smith et al. 2017). The rs-fMRI data were masked using a grey matter mask, which was derived from the MNI152 template by binarising a 75% probability grey matter mask. This restricted the analysis to voxels where CB1 and CB2 receptor expression would be expected.

The subject-specific time series estimated from the first step of the dual regression were then used as temporal regressors in the second step of the multivariate regression analysis to estimate the subject-specific spatial maps of the BOLD response after THC+CBD, THC and placebo. Both data and the design matrix were demeaned (--demean option) and the design matrix columns were normalised to unit standard deviation (--des_norm option), as is usually performed in the second stage of dual regression analyses (Filippini, MacIntosh et al. 2009). Additional technical details on dual regression implementation can be found in (Nickerson, Smith et al. 2017), and details on the REACT method in (Dipasquale, Selvaggi et al. 2019).

### Statistical analyses

The main effect of drug treatment (across placebo, THC, and THC+CBD conditions) as well as a test for a drug*age-group interaction in a voxel-wise, whole-brain manner, was carried out using a 3 × 2 mixed measures ANOVA model. F tests were used to test for significant differences between drug conditions and significant interaction effects, implemented using FSL’s ‘randomise’ module as described above.

Because the F statistics derived from the initial model are non-directional, follow-up group-level analyses were performed to calculate the directional changes in receptor-enriched functional connectivity between each drug and placebo conditions (THC+CBD>Placebo, THC- only>Placebo) for each receptor target (CB1 PET, CB1 GENE, CB2 GENE). These tests were performed using randomise (Winkler, Ridgway et al. 2014) with 5000 permutations per test and contrast, and thresholded using the Threshold-Free Cluster Enhancement (TFCE) algorithm. To compare results obtained using CB1 PET and CB1 gene maps, dice coefficients were used, calculated using custom Python code.

Additional exploratory analyses included directly testing the effects of age by comparing teens with adults. Mid-level analyses were performed for each subject to produce drug>placebo and placebo>drug images for each subject for each active compound (THC and THC+CBD). Then these outputs for the two groups were compared using randomise as described across the three receptor targets.

#### Correlations with plasma levels and subjective drug effects

Regions where there was a significant effect of drug (compared to placebo, identified by the analyses above) were used as binary masks to extract mean parameter estimates for the drug- induced changes in the CB1 and CB2 enriched connectivity for each subject (from the mid-level fixed effects analyses identifying the effect of each drug for each subject, THC+CBD>Placebo & THC-only>Placebo). These were correlated with drug plasma levels (THC for both THC+CBD and THC-only, and CBD for the THC+CBD condition) as well as with subjective drug effects using the “feel drug”, “paranoid”, “anxious” and “relaxed” visual analogue 0-10 scale scores.

To carry out exploratory voxel-wise analyses into the relationships between plasma levels and subjective drug effects, in regions where there was a significant effect of drug, Analysis of Functional Neuroimages (AFNI v.20.1.06) 3dTorr1D module was used to run voxel-wise Pearson’s correlations between receptor-enriched functional connectivity, drug plasma levels, and subjective drug effects. To identify areas of significant correlations, resultant *r*- coefficient images were transformed to *t*-statistical images using fslmaths, followed by t-maps being transformed to *Z*-scores using FSL’s ‘ttoz’ function as previously done in (Shatalina, Whitehurst et al. 2024). The Z-score images were then processed using Conn’s TFCE module (apply_tfce, E=0.5, H=2, Hmin=0). To carry out whole-brain correction consistent with the group-level randomise analyses, small clusters below 15 voxels (with connectivity set to 26) were removed using SPM’s spm_bwlabel. A Bonferroni correction was applied to the TFCE p-value. For THC and CBD levels, p<0.025 was used; for analyses of subjective drug effects, p<0.0125 was used. To provide additional visualisation of the significantly correlated clusters, the clusters were used as binary masks to extract subject-level mean values for those regions from the mid-level analyses identifying effects of each drug for each subject using FSL’s ‘fslmeants’ and plotted using ‘matplotlib’; these images are provided as visualisation references only. Regions with significant correlations were automatically identified by checking for an overlap of >10 voxels with regions of the Harvard-Oxford cortical and subcortical atlases thresholded at 25% max-probability (reported in supplementary tables 2&3).

## Results

The two-step dual regression analysis produced one subject-specific map for each receptor target and drug condition (Figure 2, showing these (unthresholded) target-enriched maps averaged across participants). We found significant decreases in both CB1 and CB2-enriched functional connectivity (p< 0.05, corrected for multiple comparisons at the cluster level using TFCE) induced by THC and THC+CBD cannabis (Figure 3, Table 2). The effect of THC+CBD on CB1-enriched functional connectivity involved a range of cortical areas, including the dorsolateral prefrontal cortex, superior frontal gyrus, cingulate cortex, insular cortex, as well as temporal regions, the hippocampus, amygdala, and the putamen. The THC+CBD induced a decrease in CB2-enriched functional connectivity in regions with a high degree of spatial overlap with the CB1-enriched maps, as shown in Figure 3. There were additional effects seen in the frontal pole, and no reductions in functional connectivity in the amygdala, putamen and DLPFC as seen in the CB1-enriched maps.

**Table 2:**

| Receptor | Regions with significant reductions in receptor-enriched functional connectivity |  |
| --- | --- | --- |
|  | THC-only | THC+CBD |
| CB1 (PET) | - Insular Cortex<br>- Amygdala<br>- Posterior Frontal Cortex<br>- Superior Frontal Gyrus<br>- Orbitofrontal Cortex<br>- DLPFC<br>- Temporal Regions | - Dorsolateral Prefrontal Cortex<br>- Superior Frontal Gyrus<br>- Cingulate Cortex<br>- Insular Cortex<br>- Temporal Regions<br>- Hippocampus<br>- Amygdala<br>- Putamen |
| CB2 (Gene) | - Superior Frontal Gyrus<br>- Postcentral Gyrus<br>- Precentral Gyrus<br>- Insular Cortex<br>- Putamen<br>- Temporal Regions | - Dorsolateral Prefrontal Cortex<br>- Superior Frontal Gyrus<br>- Cingulate Cortex<br>- Insular Cortex<br>- Temporal Regions<br>- Hippocampus<br>- Frontal Pole |

**Figure 2:**
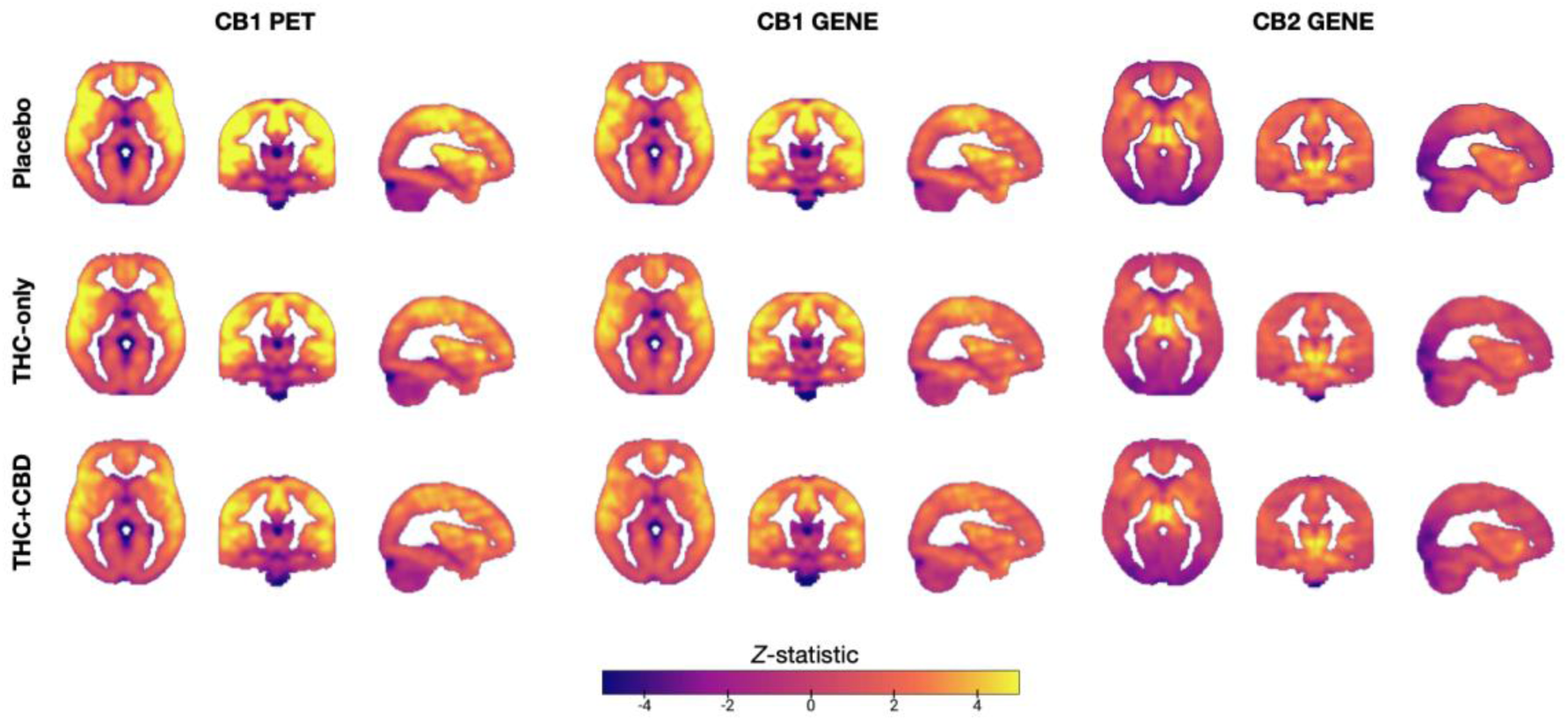
Group-mean (unthresholded) receptor-enriched functional-connectivity maps under each drug condition (n=48). REACT-derived Z-score maps for CB1 and CB2 receptors during Placebo, THC-only (8 mg/75 kg) and THC+CBD (8 mg THC + 24 mg CBD/75 kg).

**Figure 3:**
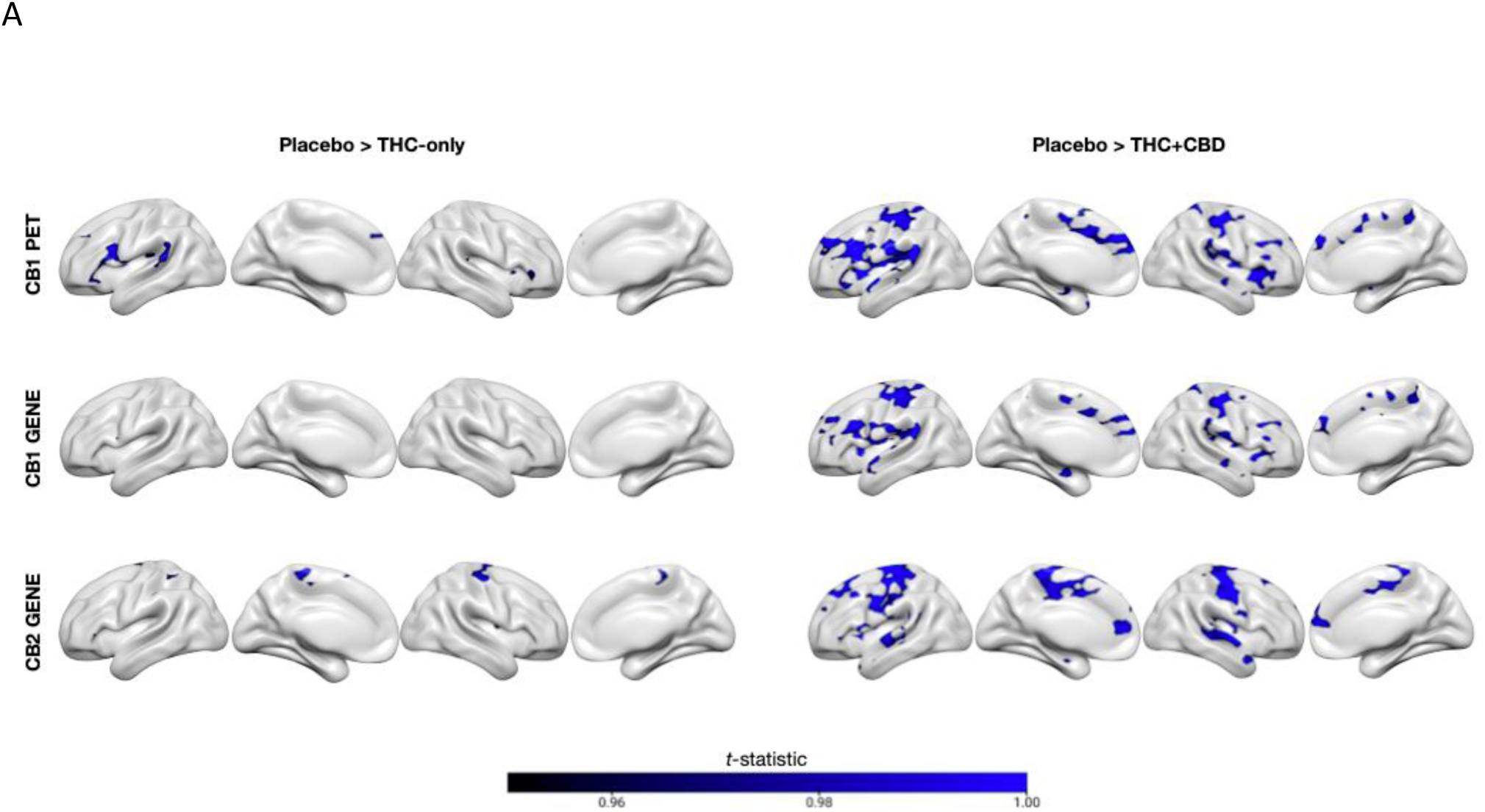
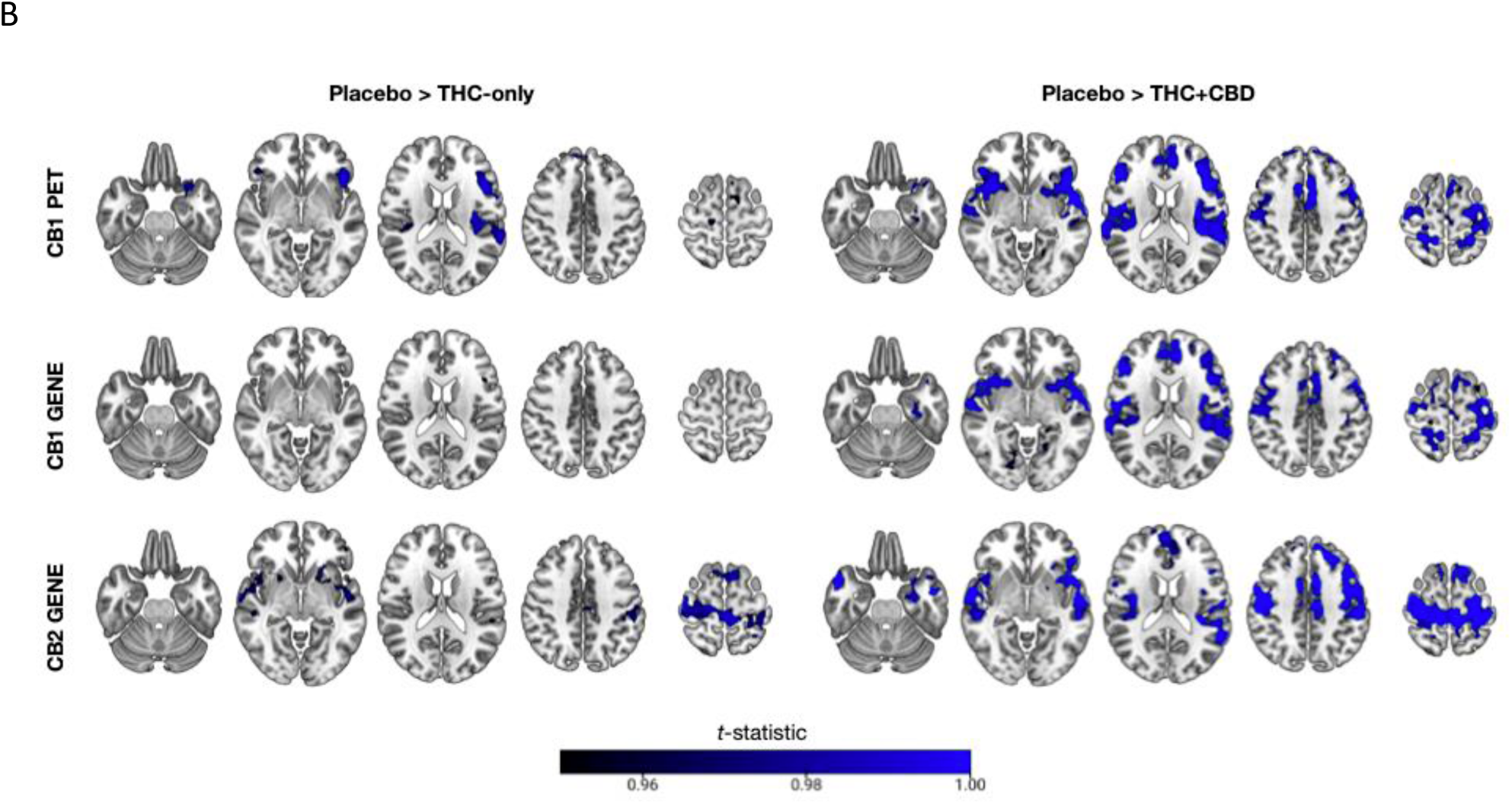
Cannabis-induced reductions in receptor-enriched connectivity (Placebo > Drug). **(A)** Surface renderings**; (B)** orthogonal slices (x = –26, –4, 18, 42, 64) showing t-statistics. Voxels showing significant decreases after THC-only or THC+CBD conditions displayed for CB1 PET, CB1 gene and CB2 gene pipelines (5000-permutation TFCE, p<0.05 FWE-corrected, n = 48).

In the THC-only condition, there were also significant functional connectivity decreases in the maps enriched by both CB1 and CB2, but these had a lesser spatial extent, although effects were seen in similar areas. For CB1, these were in the insular cortex, the amygdala, posterior frontal cortex, superior frontal gyrus, orbitofrontal cortex, DLPFC, and temporal regions. Decreases in CB2-enriched functional connectivity spanned the superior frontal gyrus, postcentral gyrus, precentral gyrus, insular cortex, putamen and temporal regions (Figure 3.

As shown in Figure 2, the CB1 gene pipeline produced similar results to the CB1 PET pipeline in the THC+CBD condition, with a dice coefficient of 0.81. In the THC-only condition, the dice coefficient was 0.07, likely due to the small number of significant voxels compared between the two pipelines (283 in the CB1-gene compared with 7137 in the CB1-PET pipelines).

However, voxels for the CB1 gene-expression enriched result were in the same regions as those for the CB1-PET enriched results.

There were no significant differences between teens and adults in any drug condition in both the CB1 and CB2 enriched functional connectivity differences, no significant differences when directly comparing the THC-only and the THC+CBD conditions, and no age*drug interactions.

We found the THC plasma concentration at 20 minutes post dosing was significantly higher in the THC+CBD visit (THC mean concentration 27.86± SD 14.20 ng/ml) compared to the THC- only visit (THC mean concentration 14.88± SD 7.15 ng/ml) (t(44) = -7.30, p < 0.0001, paired t-test). 3 subjects were missing THC plasma concentration data and were excluded from this comparison. In the THC+CBD condition, THC and CBD plasma concentrations were significantly correlated (r=0.84, p<0.001, Pearson’s correlation).

### Correlations with plasma levels

There were no significant correlations between CBD and THC plasma levels and receptor-enriched connectivity when averaged across all significant voxels for each target (supplementary table 1). In exploratory analyses in specific regions, we found that both THC and CBD plasma levels were correlated with decreases in CB1- and CB2-enriched functional connectivity induced by the administration of THC+CBD cannabis, with greater plasma levels being significantly correlated with greater decreases in connectivity (Figure 4). Significant regions included the postcentral gyrus, planum temporale, inferior frontal gyrus and insular cortex, as well as other regions (see supplementary table 2). As shown in Figure 4 below, areas where there are significant positive correlations are shown in red overlaid over the drug-induced change in receptor-enriched functional connectivity (orange), and negative correlations are shown in dark blue. THC and CBD plasma levels were also correlated with decreases in CB2-enriched functional connectivity in the precentral and post-central gyrus.

**Figure 4:**
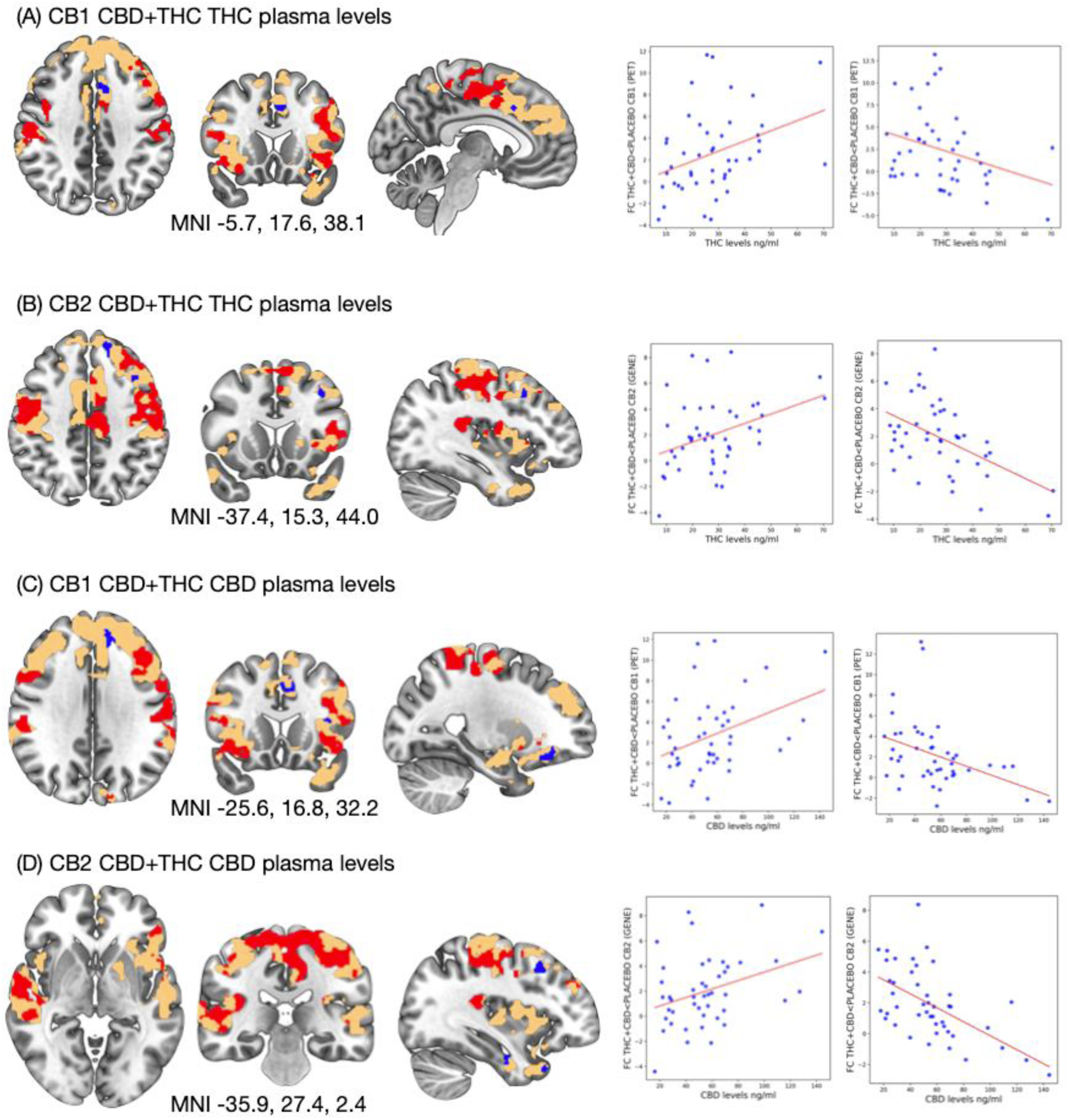
Significant relationships between plasma cannabinoid levels and the magnitude of THC+CBD-induced connectivity changes: Clusters where higher THC or CBD concentrations correlate with larger (red-yellow) or smaller (blue) decreases in CB1 and CB2-enriched connectivity (TFCE p<0.025 FWE) and corresponding scatterplots capturing average receptor- enriched connectivity changes extracted from significant clusters (included for visualisation purposes only, n = 45)

The similarity in significant correlations between both CBD and THC plasma levels can be explained by THC and CBD plasma levels being correlated (Pearson’s r=0.84, p<0.05) across individuals. There were no significant correlations between THC levels and receptor-enriched functional connectivity induced by the administration of THC-only cannabis in either the CB1 or CB2-enriched maps.

### Correlations with subjective drug effects

When data were averaged across all significant voxels for each target, there were no significant correlations between CBD and THC plasma levels and subjective drug effects (supplementary table 1). In exploratory analyses (corrected for multiple comparisons), we found that the subjective effects of the drug were both negatively and positively correlated with decreases in CB1 and CB2-enriched functional connectivity induced by both THC+CBD (figure 5, supplementary figure 1, supplementary table 3) and THC-only cannabis (supplementary figure 1, supplementary table 3). Significant associations for the ‘feel drug’ scale were identified only in the THC+CBD condition and are presented in the main text (Figure 5), with results for other subjective effect scales are shown in the supplement. Greater decreases in CB1-enriched connectivity in the opercular cortex were associated with lower feel “drug scores”, while greater decreases in CB1-enriched connectivity in the middle frontal gyrus were associated with higher “feel drug” scores (5A). Additionally, a greater decrease of CB2 enriched connectivity was associated with lower “feel drug” scores in the frontal pole, cingulate, superior frontal gyrus, as well as in the angular gyrus, temporal pole, superior and middle temporal gyri regions (5B). See supplementary table 3 for a full description of significant clusters.

**Figure 5:**
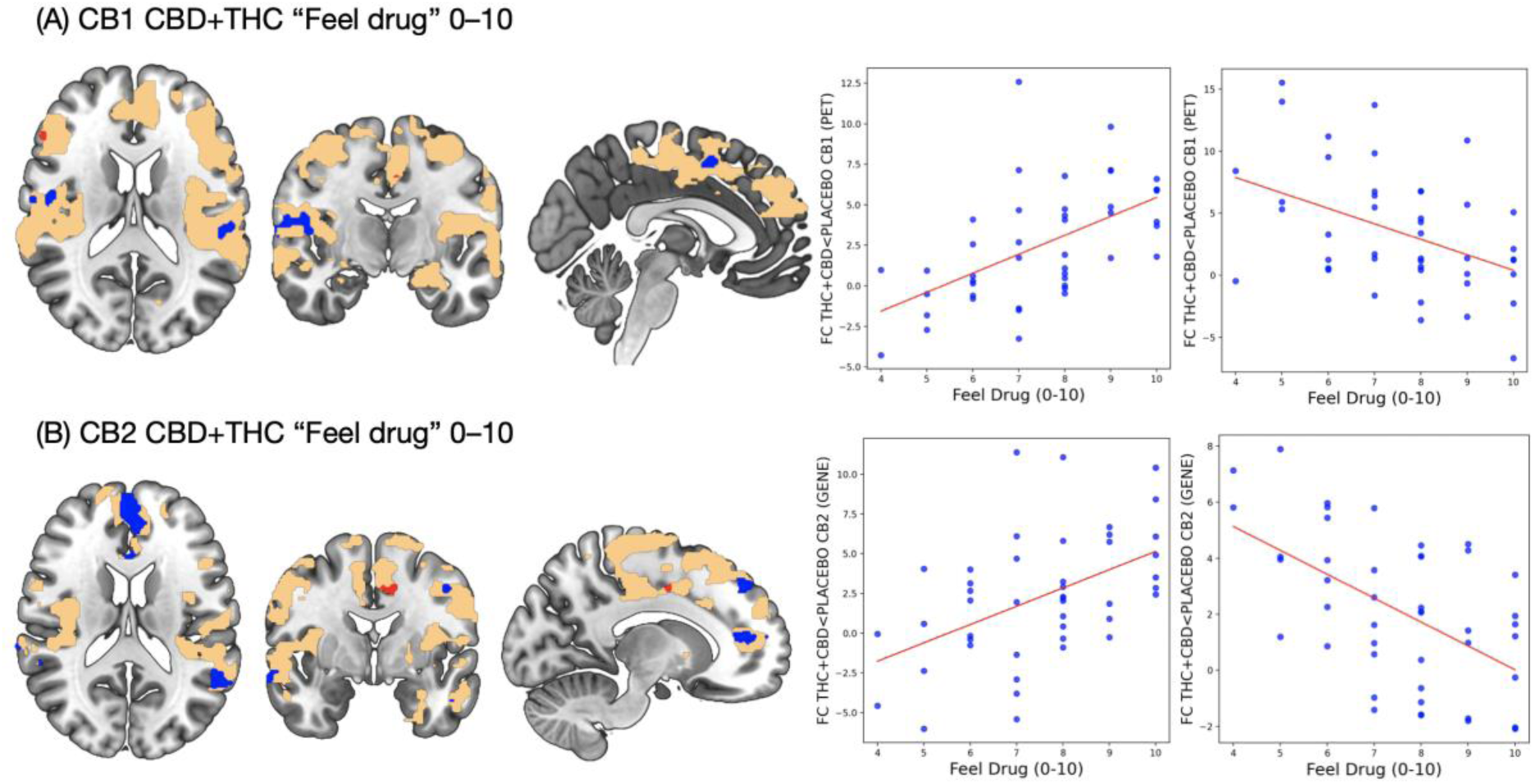
Subjective “feel drug” ratings relate to receptor-enriched connectivity after THC+CBD. Voxelwise correlations (TFCE, p<0.0125 FWE) between visual analogue “feel drug” scores and CB1 (A) and CB2 (B) enriched connectivity. Red denotes positive, blue negative associations. Scatter plots show average correlations in significant clusters and are for visualisation purposes only, n=48.

## Discussion

In our analyses, we found decreases in CB1-enriched functional connectivity in both the THC- only and the THC+CBD conditions, in regions that included the dorsolateral prefrontal cortex, superior frontal gyrus, cingulate cortex, insular cortex, as well as temporal regions, the hippocampus, amygdala, and the putamen. These results extend previous work by Ertl et al. (2024), where the authors report decreases in functional connectivity in a number of networks, including the executive control and salience networks, as well as several subcortical networks. Specifically, Ertl and al. reported decreased DLPFC and cingulate connectivity with other regions of the executive control network, decreased insula connectivity within the executive control and salience networks, decreased hippocampus-frontal and hippocampus-PCC connectivity, as well as decreased connectivity of the temporal regions within limbic and salience networks. The overlap with regions identified by REACT suggests these effects may be CB1-mediated. In addition, we identified reductions in CB1-enriched connectivity of the putamen and the amygdala, which Ertl et al did not see effects using a traditional functional connectivity approach.

We further found that CB2-enriched functional connectivity was also decreased following both the administration of THC-only and THC+CBD compared to placebo, spanning similar regions to changes observed in CB1-enriched functional connectivity. In the THC+CBD treatment, additional effects were seen in the frontal pole, with no reductions in functional connectivity in the amygdala, putamen and DLPFC that were seen with the CB1-enriched maps. These results extend prior work by indicating that functional connectivity decreases in different regions may be mediated via different receptors.

We found that THC+CBD cannabis caused decreases in functional connectivity that had a greater spatial extent than in the THC-only treatment, in both CB1 and CB2-enriched functional connectivity. This is consistent with results reported by Ertl et al., where THC+CBD similarly induced greater decreases in functional connectivity, but in opposition to Wall et al. (2019), where the authors found THC-only treatment caused greater decreases in functional connectivity than THC co-administered with CBD (Wall, Pope et al. 2019). We also saw that THC plasma levels were significantly higher in the THC+CBD condition compared to the THC- only condition, despite the same controlled method of administration. As discussed by Ertl et al., this may be due to CBD and THC competing for metabolism at CYP450 enzymes, which results in significantly higher THC plasma levels when it is co-administered with CBD (Arkell, Lintzeris et al. 2019, Zamarripa, Spindle et al. 2023, Ertl, Freeman et al. 2024). Other recent work has provided additional evidence for this, showing that co-administration of 450mg of CBD with 9mg of THC significantly increased subjective and behavioural effects of THC, along with increases in blood-plasma levels and elimination half-life of THC (Gorbenko, Heuberger et al.), and that THC co-administered with CBD resulted in higher THC levels than when administered alone (Arkell, Lintzeris et al. 2019). Thus, it is most likely that we are observing a THC dose-response effect across both receptors. Supporting this interpretation, in the THC+CBD condition, plasma levels of both THC and CBD were correlated with decreases in CB1-enriched functional connectivity in regions including the postcentral gyrus, planum temporale, inferior frontal gyrus, and insular cortex, as well as with decreases in CB2-enriched connectivity in the precentral and postcentral gyrus. Since THC and CBD plasma levels were correlated across subjects due to co-administration, it is difficult to test which one may be driving the positive relationship in these regions and indicate future studies would benefit from a CBD-only condition.

Taken together, these results are consistent with greater THC dose driving greater decreases in receptor-enriched functional connectivity. However, we did not observe a similar correlation between THC levels and receptor-enriched connectivity in the THC-only condition. One explanation for this is the smaller number of voxels included in the analysis, as we only analysed voxels showing significant drug (THC>placebo) effects (see figure 2). Thus, following cluster-correction, there is a very low likelihood that there would be significant overlapping clusters, additionally, lower variation in THC plasma levels in the THC-only condition could further reduce our ability to detect an association.

We found that subjective drug effects were related to both CB1 and CB2-enriched functional connectivity in a region-specific manner in the CBD+THC condition only. Similarly to analyses with THC and CBD levels, correlations were performed only in regions where there was a significant effect of the drug. Thus, this may explain why there were no effects in the THC- only condition, which was also associated with lower THC levels. In exploratory analyses, we found that greater decreases in CB1-enriched connectivity in the opercular cortex were associated with lower feel “drug scores”, while greater decreases in CB1-enriched connectivity in the middle frontal gyrus were associated with higher “feel drug” scores. One interpretation of the negative correlation with interoceptive/sensorimotor regions is that CB1 activations may be linked to lower awareness of the effect of the drug, resulting in a lower “feel drug” rating. Additionally, a greater decrease of CB2 enriched connectivity was associated with lower “feel drug” scores in the frontal pole, cingulate, superior frontal gyrus, as well as in the angular gyrus, temporal pole, superior and middle temporal gyri regions. This raises questions around the roles CB2 may play in mediating the subjective effects of cannabis.

Previous fMRI studies have linked THC-induced changes in perceived euphoria/high states with altered connectivity, specifically with increases in frontostriatal connectivity (Van Hell, Bossong et al. 2011) and PCC-sensorimotor/precentral gyrus connectivity (Kleinloog, Rombouts et al. 2015). In opposition to this, a study co-administering CBD with THC found ratings of how “high” participants felt were related to greater reductions in PCC connectivity (Wall, Pope et al. 2019). Our findings add to this body of literature by identifying associations with brain function related to specific receptor systems.

### Strengths and limitations

This data set has a number of strengths, including groups of adolescents and adults with an equal split of males and females, a relatively large sample size compared to previous studies of this type (Winton-Brown, Allen et al. 2011, Klumpers, Cole et al. 2012, Bossong, van Hell et al. 2013, Wall, Pope et al. 2019), and a well-matched placebo control treatment. The two groups were also well-matched in terms of cannabis use frequency. In addition, vaporised cannabis ensures better absorption and ecological validity than oral THC and CBD challenges, avoiding first-pass metabolism and achieving a relevant concentration of CBD to achieve negative-allosteric modulation of CB1 (Lawn, Trinci et al. 2023).

A key consideration for this study is the absence of a PET tracer for CB2 receptors, which led us to use the Allen Human Brain Atlas (AHBA) to define CB2-mediated changes in functional connectivity. Previous validation studies have demonstrated good concordance between PET imaging and AHBA maps for receptors that do not undergo significant post-translational modifications (Rizzo, Veronese et al. 2014, Selvaggi, Rizzo et al. 2021, Hansen, Markello et al. 2022). Post-translational modifications, such as phosphorylation, glycosylation, or ubiquitination, can alter the structure, function, or localisation of receptor proteins, potentially affecting their availability and binding properties. As receptors like CB1 and CB2 do not undergo substantial post-translational modifications (Carruthers and Grimsey 2024), they are less likely to show variability in receptor expression or distribution due to these molecular changes, supporting the use of AHBA data to represent CB2 receptor distribution in this study. We additionally included an analysis of CB1-mediated changes using the AHBA to define CB1-receptor distribution. A comparison of the CB1 AHBA and CB1 PET REACT pipelines showed a high degree of spatial overlap in the results, further suggesting AHBA data is suitable for these analyses. However, it must be cautioned that AHBA data is derived from 6 subjects only, and has limited spatial sampling, and thus, results should be interpreted with some caution.

Another key consideration is that this study included participants who were (semi-)regular cannabis users, while the PET/AHBA reference maps were acquired from averages of normal “healthy” subjects. Repeated administration of receptor agonists may result in receptor internalisation and/or a reduction in receptor protein signalling (Pertwee 2006) and significant CB1 density alteration has been seen in daily cannabis users (Ceccarini, Kuepper et al. 2015). However, given that participants in this study used cannabis 1.5 days/week on average, it is likely that any receptor alterations would be relatively minimal and the ‘normal’ spatial distribution of receptors would be preserved. Replicating these results in a multimodal PET-fMRI study where individual receptor distribution data are available for each subject would answer to what extent individual variability in receptor distribution affects REACT results.

Finally, it is important to note that the co-administration of CBD+THC resulted in higher plasma THC levels than in the THC-only condition, meaning that in the THC+CBD condition, two key variables changed, affecting the ability to make a clear interpretation of the results. Results from previous studies are mixed and conclude that CBD may be both oppositional to THC (Winton-Brown, Allen et al. 2011), enhance its effects (Gorbenko, Heuberger et al.), or have no effects at all (Englund, Oliver et al. 2023). Overall, while the increase in THC plasma level we saw confounds the pharmacological interpretation of the study, the comparison is still instructive as it effectively mimics what happens in real-world cannabis use.

### Implications and conclusions

The results show that cannabinoids reliably decrease resting-state activity in a broad network of regions associated with the distribution maps of both CB1 and CB2 receptors and suggest that co-administration of CBD with THC enhances these effects. This is evidence that cannabis with high CBD levels may not necessarily attenuate the neural effects of THC (and may in fact produce stronger effects). The negative finding of no interaction between cannabis treatments and age group suggests that effects on resting-state activity related to cannabinoid receptor distribution do not significantly differ between adolescents and adults. This indicates that adolescents do not appear to be more at risk for negative effects than adults. Further work in this area using multi-modal imaging (PET with suitable cannabinoid ligands, and fMRI) in the same cohort of subjects would be highly valuable and would further advance our understanding of the effects of cannabinoids on brain function and physiology.

### Financial disclosures

MBW and NE’s primary employer is Perceptive LLC, a contract research organisation that performs commercial research for the pharmaceutical and biotechnology industries. OH is a part-time employee of H Lundbeck A/s. He has received investigator-initiated research funding from and/or participated in advisory/ speaker meetings organised by Angellini, Autifony, Biogen, Boehringer-Ingelheim, Eli Lilly, Heptares, Global Medical Education, Invicro, Janssen, Lundbeck, Neurocrine, Otsuka, Sunovion, Recordati, Roche and Viatris/ Mylan. HVC has consulted for Janssen Research and Development. The remaining authors declare no competing interests.

## References

1. Arkell, T. R., N. Lintzeris, R. C. Kevin, J. G. Ramaekers, R. Vandrey, C. Irwin, P. S. Haber and I. S. McGregor (2019). “Cannabidiol (CBD) content in vaporized cannabis does not prevent tetrahydrocannabinol (THC)-induced impairment of driving and cognition.” Psychopharmacology 236(9): 2713–2724.

2. Atwood, B. K., A. Straiker and K. Mackie (2012). “CB2 cannabinoid receptors inhibit synaptic transmission when expressed in cultured autaptic neurons.” Neuropharmacology 63(4): 514–523.

3. Bossong, M. G., H. H. van Hell, G. Jager, R. S. Kahn, N. F. Ramsey and J. M. Jansma (2013). “The endocannabinoid system and emotional processing: a pharmacological fMRI study withΔ 9-tetrahydrocannabinol.” European Neuropsychopharmacology 23(12): 1687–1697.

4. Carruthers, E. R. and N. L. Grimsey (2024). “Cannabinoid CB2 receptor orthologues; in vitro function and perspectives for preclinical to clinical translation.” British Journal of Pharmacology 181(14): 2247–2269.

5. Ceccarini, J., R. Kuepper, D. Kemels, J. van Os, C. Henquet and K. Van Laere (2015). “[18 F] MK-9470 PET measurement of cannabinoid CB 1 receptor availability in chronic cannabis users.” Addiction biology 20(2): 357–367.

6. Curran, H. V., T. P. Freeman, C. Mokrysz, D. A. Lewis, C. J. Morgan and L. H. Parsons (2016). “Keep off the grass? Cannabis, cognition and addiction.” Nature Reviews Neuroscience 17(5): 293–306.

7. Demetriou, L., O. S. Kowalczyk, G. Tyson, T. Bello, R. D. Newbould and M. B. Wall (2018). “A comprehensive evaluation of increasing temporal resolution with multiband-accelerated protocols and effects on statistical outcome measures in fMRI.” Neuroimage 176: 404–416.

8. den Boon, F. S., P. Chameau, Q. Schaafsma-Zhao, W. van Aken, M. Bari, S. Oddi, C. G. Kruse, M. Maccarrone, W. J. Wadman and T. R. Werkman (2012). “Excitability of prefrontal cortical pyramidal neurons is modulated by activation of intracellular type-2 cannabinoid receptors.” Proceedings of the National Academy of Sciences 109(9): 3534–3539.

9. Dipasquale, O., D. Martins, A. Sethi, M. Veronese, S. Hesse, M. Rullmann, O. Sabri, F. Turkheimer, N. A. Harrison and M. A. Mehta (2020). “Unravelling the effects of methylphenidate on the dopaminergic and noradrenergic functional circuits.” Neuropsychopharmacology 45(9): 1482–1489.

10. Dipasquale, O., P. Selvaggi, M. Veronese, A. S. Gabay, F. Turkheimer and M. A. Mehta (2019). “Receptor-Enriched Analysis of functional connectivity by targets (REACT): A novel, multimodal analytical approach informed by PET to study the pharmacodynamic response of the brain under MDMA.” Neuroimage 195: 252–260.

11. Dukart, J., S. Holiga, M. Rullmann, R. Lanzenberger, P. C. Hawkins, M. A. Mehta, S. Hesse, H. Barthel, O. Sabri and R. Jech (2021). JuSpace: A tool for spatial correlation analyses of magnetic resonance imaging data with nuclear imaging derived neurotransmitter maps, Wiley Online Library.

12. Elmes, S. J., M. D. Jhaveri, D. Smart, D. A. Kendall and V. Chapman (2004). “Cannabinoid CB2 receptor activation inhibits mechanically evoked responses of wide dynamic range dorsal horn neurons in naive rats and in rat models of inflammatory and neuropathic pain.” European Journal of Neuroscience 20(9): 2311–2320.

13. Englund, A., D. Oliver, E. Chesney, L. Chester, J. Wilson, S. Sovi, A. De Micheli, J. Hodsoll, P. Fusar-Poli and J. Strang (2023). “Does cannabidiol make cannabis safer? A randomised, double-blind, cross-over trial of cannabis with four different CBD: THC ratios.” Neuropsychopharmacology 48(6): 869–876.

14. Ertl, N., T. P. Freeman, C. Mokrysz, S. Ofori, A. Borissova, K. Petrilli, H. V. Curran, W. Lawn and M. B. Wall (2024). “Acute effects of different types of cannabis on young adult and adolescent resting-state brain networks.” Neuropsychopharmacology: 1–12.

15. Filippini, N., B. J. MacIntosh, M. G. Hough, G. M. Goodwin, G. B. Frisoni, S. M. Smith, P. M. Matthews, C. F. Beckmann and C. E. Mackay (2009). “Distinct patterns of brain activity in young carriers of the APOE-ε4 allele.” Proceedings of the National Academy of Sciences 106(17): 7209–7214.

16. Franklin, J. M. and G. A. Carrasco (2012). “Cannabinoid-induced enhanced interaction and protein levels of serotonin 5-HT2A and dopamine D2 receptors in rat prefrontal cortex.” Journal of psychopharmacology 26(10): 1333–1347.

17. Freeman, A. M., K. Petrilli, R. Lees, C. Hindocha, C. Mokrysz, H. V. Curran, R. Saunders and T. P. Freeman (2019). “How does cannabidiol (CBD) influence the acute effects of delta-9-tetrahydrocannabinol (THC) in humans? A systematic review.” Neuroscience & Biobehavioral Reviews 107: 696–712.

18. Freeman, T. P. and V. Lorenzetti (2020). “‘Standard THC units’: a proposal to standardize dose across all cannabis products and methods of administration.” Addiction 115(7): 1207–1216.

19. Gorbenko, A. A., J. A. Heuberger, L. E. Klumpers, M. L. de Kam, P. K. Strugala, S. J. de Visser and G. J. Groeneveld “Cannabidiol Increases Psychotropic Effects and Plasma Concentrations of Δ9-Tetrahydrocannabinol Without Improving Its Analgesic Properties.” Clinical Pharmacology & Therapeutics.

20. Grabon, W., S. Rheims, J. Smith, J. Bodennec, A. Belmeguenai and L. Bezin (2023). “CB2 receptor in the CNS: From immune and neuronal modulation to behavior.” Neuroscience & Biobehavioral Reviews 150: 105226.

21. Hansen, J. Y., R. D. Markello, L. Tuominen, M. Nørgaard, E. Kuzmin, N. Palomero-Gallagher, A. Dagher and B. Misic (2022). “Correspondence between gene expression and neurotransmitter receptor and transporter density in the human brain.” NeuroImage 264: 119671.

22. Hansen, J. Y., G. Shafiei, R. D. Markello, K. Smart, S. M. Cox, M. Nørgaard, V. Beliveau, Y. Wu, J.-D. Gallezot and É. Aumont (2022). “Mapping neurotransmitter systems to the structural and functional organization of the human neocortex.” Nature neuroscience 25(11): 1569–1581.

23. Hwang, E.-S., H.-B. Kim, S. Lee, M.-J. Kim, K.-J. Kim, G. Han, S.-Y. Han, E.-A. Lee, J.-H. Yoon and D.-O. Kim (2020). “Antidepressant-like effects of β-caryophyllene on restraint plus stress-induced depression.” Behavioural brain research 380: 112439.

24. Jhaveri, M., S. Elmes, D. Richardson, D. Barrett, D. Kendall, R. Mason and V. Chapman (2008). “Evidence for a novel functional role of cannabinoid CB2 receptors in the thalamus of neuropathic rats.” European Journal of Neuroscience 27(7): 1722–1730.

25. Kano, M., T. Ohno-Shosaku, Y. Hashimotodani, M. Uchigashima and M. Watanabe (2009). “Endocannabinoid-mediated control of synaptic transmission.” Physiological reviews.

26. Kleinloog, D., S. Rombouts, R. Zoethout, L. Klumpers, M. Niesters, N. Khalili-Mahani, A. Dahan and J. van Gerven (2015). “Subjective Effects of Ethanol, Morphine, Δ9- Tetrahydrocannabinol, and Ketamine Following a Pharmacological Challenge Are Related to Functional Brain Connectivity.” Brain connectivity 5(10): 641–648.

27. Klumpers, L. E., D. M. Cole, N. Khalili-Mahani, R. P. Soeter, E. T. Te Beek, S. A. Rombouts and J. M. van Gerven (2012). “Manipulating brain connectivity with δ9- tetrahydrocannabinol: a pharmacological resting state FMRI study.” Neuroimage 63(3): 1701–1711.

28. Laprairie, R., A. Bagher, M. Kelly and E. Denovan-Wright (2015). “Cannabidiol is a negative allosteric modulator of the cannabinoid CB1 receptor.” British journal of pharmacology 172(20): 4790–4805.

29. Lawn, T., O. Dipasquale, A. Vamvakas, I. Tsougos, M. A. Mehta and M. A. Howard (2022). “Differential contributions of serotonergic and dopaminergic functional connectivity to the phenomenology of LSD.” Psychopharmacology 239(6): 1797–1808.

30. Lawn, W., K. Trinci, C. Mokrysz, A. Borissova, S. Ofori, K. Petrilli, M. Bloomfield, Z. R. Haniff, D. Hall and N. Fernandez-Vinson (2023). “The acute effects of cannabis with and without cannabidiol in adults and adolescents: a randomised, double-blind, placebo-controlled, crossover experiment.” Addiction 118(7): 1282–1294.

31. Ma, Z., F. Gao, B. Larsen, M. Gao, Z. Luo, D. Chen, X. Ma, S. Qiu, Y. Zhou and J. Xie (2019). “Mechanisms of cannabinoid CB2 receptor-mediated reduction of dopamine neuronal excitability in mouse ventral tegmental area.” EBioMedicine 42: 225–237.

32. Mackie, K. (2008). “Cannabinoid receptors: where they are and what they do.” Journal of neuroendocrinology 20: 10–14.

33. Mokrysz, C., T. P. Freeman, S. Korkki, K. Griffiths and H. V. Curran (2016). “Are adolescents more vulnerable to the harmful effects of cannabis than adults? A placebo-controlled study in human males.” Translational psychiatry 6(11): e961–e961.

34. Mokrysz, C., N. D. Shaban, T. P. Freeman, W. Lawn, R. A. Pope, C. Hindocha, A. Freeman, M. B. Wall, M. A. Bloomfield and C. J. Morgan (2021). “Acute effects of cannabis on speech illusions and psychotic-like symptoms: two studies testing the moderating effects of cannabidiol and adolescence.” Psychological Medicine 51(12): 2134–2142.

35. Morgan, N. H., I. M. Stanford and G. L. Woodhall (2009). “Functional CB2 type cannabinoid receptors at CNS synapses.” Neuropharmacology 57(4): 356–368.

36. Nackley, A. G., A. M. Zvonok, A. Makriyannis and A. G. Hohmann (2004). “Activation of cannabinoid CB2 receptors suppresses C-fiber responses and windup in spinal wide dynamic range neurons in the absence and presence of inflammation.” Journal of neurophysiology 92(6): 3562–3574.

37. Nickerson, L. D., S. M. Smith, D. Öngür and C. F. Beckmann (2017). “Using dual regression to investigate network shape and amplitude in functional connectivity analyses.” Frontiers in neuroscience 11: 231426.

38. Normandin, M. D., M.-Q. Zheng, K.-S. Lin, N. S. Mason, S.-F. Lin, J. Ropchan, D. Labaree, S. Henry, W. A. Williams and R. E. Carson (2015). “Imaging the cannabinoid cb1 receptor in humans with [11c] omar: assessment of kinetic analysis methods, test– retest reproducibility, and gender differences.” Journal of Cerebral Blood Flow & Metabolism 35(8): 1313–1322.

39. Pelgrim, T. A., J. G. Ramaekers, M. B. Wall, T. P. Freeman and M. G. Bossong (2023). “Acute effects of Δ9-tetrahydrocannabinol (THC) on resting state connectivity networks and impact of COMT genotype: A multi-site pharmacological fMRI study.” Drug and alcohol dependence 251: 110925.

40. Pertwee, R. (2008). “The diverse CB1 and CB2 receptor pharmacology of three plant cannabinoids: Δ9-tetrahydrocannabinol, cannabidiol and Δ9-tetrahydrocannabivarin.” British journal of pharmacology 153(2): 199–215.

41. Pertwee, R. G. (2006). “The pharmacology of cannabinoid receptors and their ligands: an overview.” International journal of obesity 30(1): S13–S18.

42. Rizzo, G., M. Veronese, R. A. Heckemann, S. Selvaraj, O. D. Howes, A. Hammers, F. E. Turkheimer and A. Bertoldo (2014). “The predictive power of brain mRNA mappings for in vivo protein density: a positron emission tomography correlation study.” Journal of Cerebral Blood Flow & Metabolism 34(5): 827–835.

43. Sadanandan, S. M., T. Kreko-Pierce, S. N. Khatri and J. R. Pugh (2020). “Cannabinoid type 2 receptors inhibit GABAA receptor-mediated currents in cerebellar Purkinje cells of juvenile mice.” PLoS One 15(5): e0233020.

44. Selvaggi, P., G. Rizzo, M. A. Mehta, F. E. Turkheimer and M. Veronese (2021). “Integration of human whole-brain transcriptome and neuroimaging data: Practical considerations of current available methods.” Journal of neuroscience methods 355: 109128.

45. Shatalina, E., T. S. Whitehurst, E. C. Onwordi, B. J. Gilbert, G. Rizzo, A. Whittington, A. Mansur, H. Tsukada, T. R. Marques and S. Natesan (2024). “Mitochondrial complex I density is associated with IQ and cognition in cognitively healthy adults: an in vivo [18F] BCPP-EF PET study.” EJNMMI research 14(1): 1–10.

46. Shen, E. H., C. C. Overly and A. R. Jones (2012). “The Allen Human Brain Atlas: comprehensive gene expression mapping of the human brain.” Trends in neurosciences 35(12): 711–714.

47. Smith, D. V., A. V. Utevsky, A. R. Bland, N. Clement, J. A. Clithero, A. E. Harsch, R. M. Carter and S. A. Huettel (2014). “Characterizing individual differences in functional connectivity using dual-regression and seed-based approaches.” Neuroimage 95: 1–12.

48. Sokal, D., S. Elmes, D. Kendall and V. Chapman (2003). “Intraplantar injection of anandamide inhibits mechanically-evoked responses of spinal neurones via activation of CB2 receptors in anaesthetised rats.” Neuropharmacology 45(3): 404–411.

49. Spindle, T. R., E. J. Cone, N. J. Schlienz, J. M. Mitchell, G. E. Bigelow, R. Flegel, E. Hayes and R. Vandrey (2018). “Acute effects of smoked and vaporized cannabis in healthy adults who infrequently use cannabis: a crossover trial.” JAMA network open 1(7): e184841–e184841.

50. Stempel, A. V., A. Stumpf, H.-Y. Zhang, T. Özdoğan, U. Pannasch, A.-K. Theis, D.-M. Otte, A. Wojtalla, I. Rácz and A. Ponomarenko (2016). “Cannabinoid type 2 receptors mediate a cell type-specific plasticity in the hippocampus.” Neuron 90(4): 795–809.

51. Van Hell, H. H., M. G. Bossong, G. Jager, G. Kristo, M. J. Van Osch, F. Zelaya, R. S. Kahn and N. F. Ramsey (2011). “Evidence for involvement of the insula in the psychotropic effects of THC in humans: a double-blind, randomized pharmacological MRI study.” International Journal of Neuropsychopharmacology 14(10): 1377–1388.

52. Wall, M. B., R. Pope, T. P. Freeman, O. S. Kowalczyk, L. Demetriou, C. Mokrysz, C. Hindocha, W. Lawn, M. A. Bloomfield and A. M. Freeman (2019). “Dissociable effects of cannabis with and without cannabidiol on the human brain’s resting-state functional connectivity.” Journal of Psychopharmacology 33(7): 822–830.

53. White, C. M. (2019). “A review of human studies assessing cannabidiol’s (CBD) therapeutic actions and potential.” The Journal of Clinical Pharmacology 59(7): 923–934.

54. Winkler, A. M., G. R. Ridgway, M. A. Webster, S. M. Smith and T. E. Nichols (2014). “Permutation inference for the general linear model.” Neuroimage 92: 381–397.

55. Winton-Brown, T. T., P. Allen, S. Bhattacharrya, S. J. Borgwardt, P. Fusar-Poli, J. A. Crippa, M. L. Seal, R. Martin-Santos, D. Ffytche and A. W. Zuardi (2011). “Modulation of auditory and visual processing by delta-9-tetrahydrocannabinol and cannabidiol: an FMRI study.” Neuropsychopharmacology 36(7): 1340–1348.

56. Yu, H., X. Liu, B. Chen, C. R. Vickstrom, V. Friedman, T. J. Kelly, X. Bai, L. Zhao, C. J. Hillard and Q.-S. Liu (2021). “The neuroprotective effects of the CB2 agonist GW842166x in the 6-OHDA mouse model of Parkinson’s disease.” Cells 10(12): 3548.

57. Zamarripa, C. A., T. R. Spindle, R. Surujunarain, E. M. Weerts, S. Bansal, J. D. Unadkat, M. F. Paine and R. Vandrey (2023). “Assessment of orally administered Δ9- tetrahydrocannabinol when coadministered with cannabidiol on Δ9- tetrahydrocannabinol pharmacokinetics and pharmacodynamics in healthy adults: A randomized clinical trial.” JAMA Network Open 6(2): e2254752–e2254752.

58. Zhang, H.-Y., M. Gao, Q.-R. Liu, G.-H. Bi, X. Li, H.-J. Yang, E. L. Gardner, J. Wu and Z.-X. Xi (2014). “Cannabinoid CB2 receptors modulate midbrain dopamine neuronal activity and dopamine-related behavior in mice.” Proceedings of the National Academy of Sciences 111(46): E5007–E5015.

59. Zhang, H.-Y., H. Shen, M. Gao, Z. Ma, B. J. Hempel, G.-H. Bi, E. L. Gardner, J. Wu and Z.- X. Xi (2021). “Cannabinoid CB2 receptors are expressed in glutamate neurons in the red nucleus and functionally modulate motor behavior in mice.” Neuropharmacology 189: 108538.

60. Zhang, H. Y., M. Gao, H. Shen, G. H. Bi, H. J. Yang, Q. R. Liu, J. Wu, E. L. Gardner, A. Bonci and Z. X. Xi (2017). “Expression of functional cannabinoid CB2 receptor in VTA dopamine neurons in rats.” Addiction biology 22(3): 752–765.

