## Supplementary materials for "Receptor-enriched analysis of functional connectivity (REACT) for understanding cannabinoid neuropsychopharmacology"

**Supplementary Table 1:** Correlations between average receptor-enriched functional connectivity (FC) and cannabinoid plasma levels or subjective measures

| Condition | Variable of interest | Average receptor enriched functional connectivity | Correlation Coefficient (r) | p-value |
|---|---|---|---|---|
| THC +CBD | THC levels | THC+CBD<PLACEBO CB1 (PET) | 0.207787 | 0.170795 |
| THC +CBD | THC levels | THC+CBD<PLACEBO CB1 (GENE) | 0.183540 | 0.227487 |
| THC +CBD | THC levels | THC+CBD<PLACEBO CB2 (GENE) | 0.188961 | 0.21381 |
| THC +CBD | CBD levels | THC+CBD<PLACEBO CB1 (PET) | 0.228183 | 0.131658 |
| THC +CBD | CBD levels | THC+CBD<PLACEBO CB1 (GENE) | 0.190441 | 0.210179 |
| THC +CBD | CBD levels | THC+CBD<PLACEBO CB2 (GENE) | 0.149291 | 0.327682 |
| THC-only | THC levels | THC+CBD<PLACEBO CB1 (PET) | 0.010094 | 0.94753 |
| THC-only | THC levels | THC+CBD<PLACEBO CB1 (GENE) | -0.022573 | 0.882988 |
| THC-only | THC levels | THC+CBD<PLACEBO CB2 (GENE) | 0.122255 | 0.423688 |
| THC+CBD | feel drug (0-10) | THC+CBD<PLACEBO CB1 (PET) | -0.020723 | 0.892518 |
| THC+CBD | feel drug (0-10) | THC+CBD<PLACEBO CB1 (GENE) | 0.017331 | 0.910035 |
| THC+CBD | feel drug (0-10) | THC+CBD<PLACEBO CB2 (GENE) | -0.064929 | 0.67175 |
| THC-only | feel drug (0-10) | THC+CBD<PLACEBO CB1 (PET) | -0.027762 | 0.856349 |
| THC-only | feel drug (0-10) | THC+CBD<PLACEBO CB1 (GENE) | -0.174114 | 0.252669 |
| THC-only | feel drug (0-10) | THC+CBD<PLACEBO CB2 (GENE) | 0.010565 | 0.945085 |

**Supplementary Figure 1**

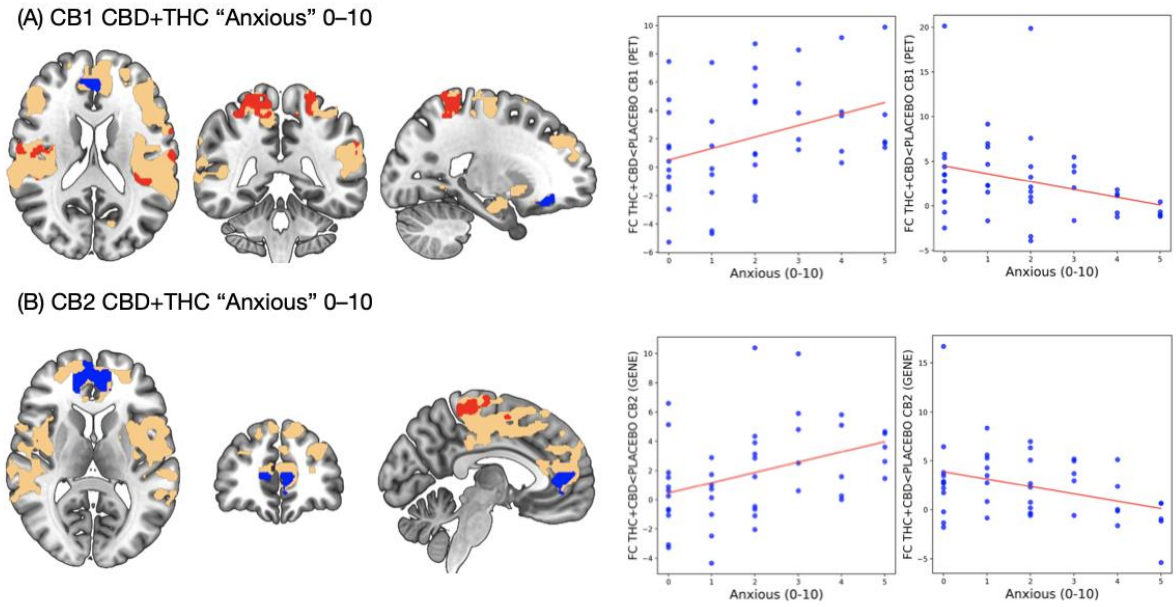

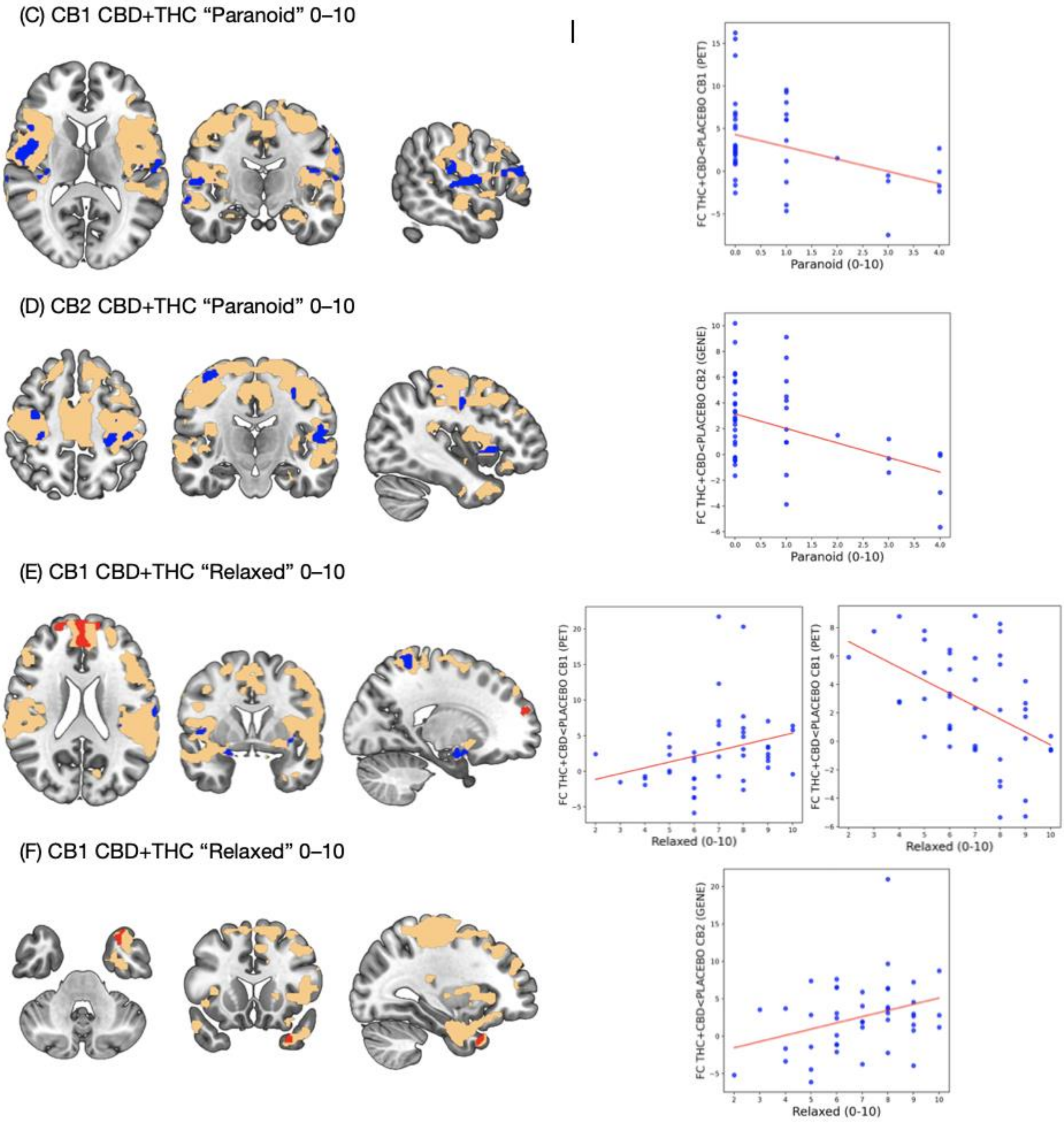


**Supplementary figure 1: Subjective drug effects ratings relate to receptor-enriched connectivity in the THC+CBD condition (drug>placebo).** Voxelwise correlations (TFCE, $p<0.0125$ FWE) between visual analogue “anxious” (A&B), “paranoid” (C&D) and “relaxed” (E&F) scores and CB1 and CB2 enriched connectivity. Red denotes positive, blue negative associations. Scatter plots show average correlations in clusters identified as significant and are for visualisation purposes only.

**Supplementary Figure 2**

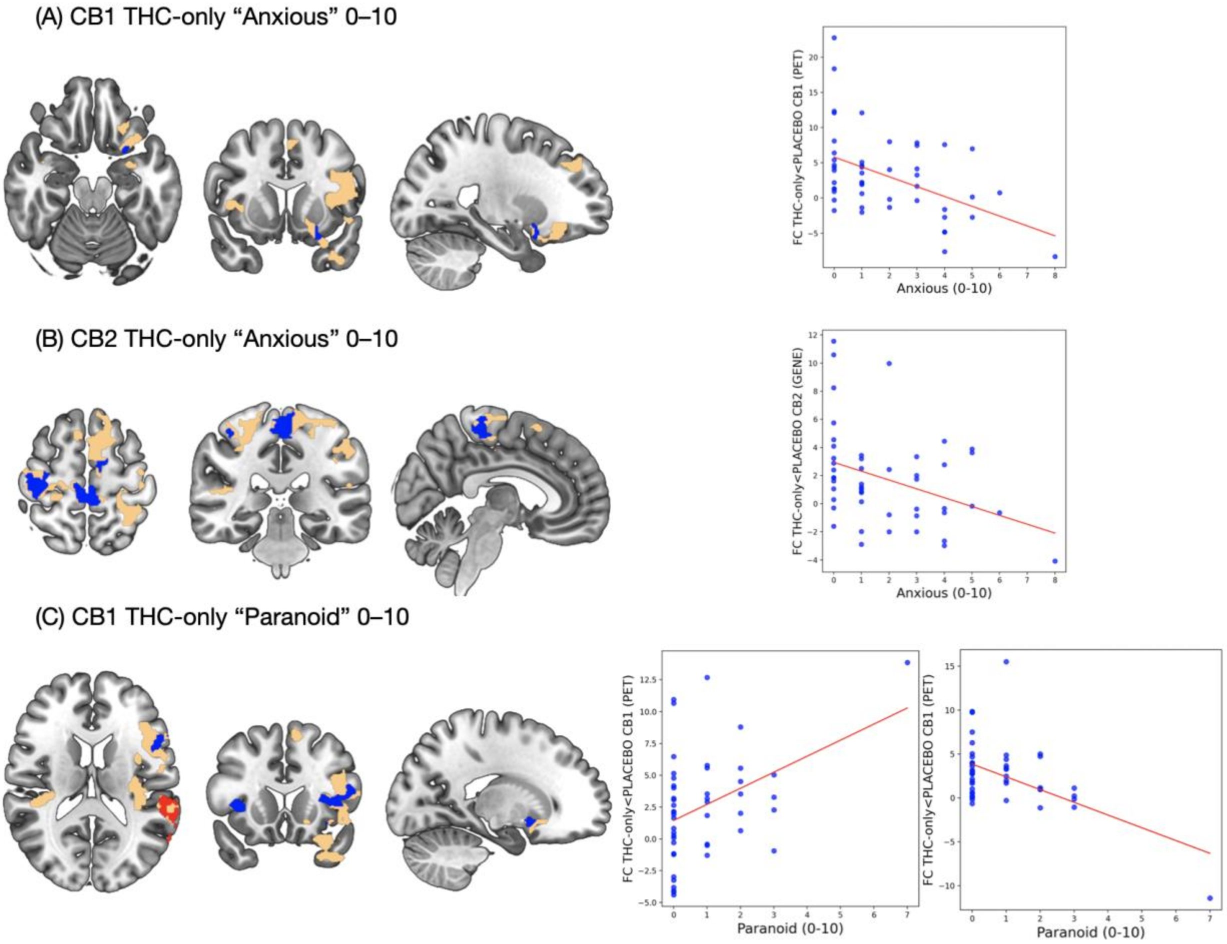

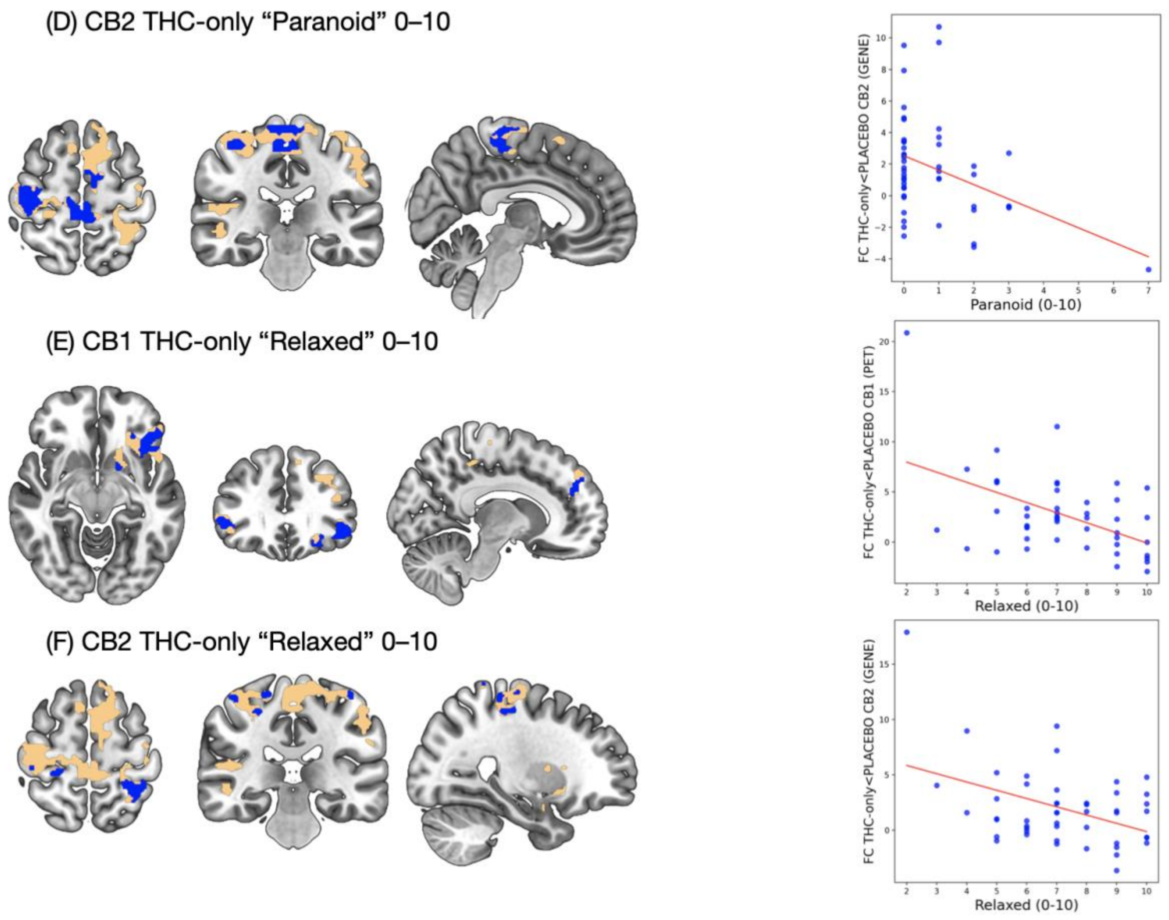


**Supplementary figure 2: Subjective drug effects ratings relate to receptor-enriched connectivity in the THC+only condition (drug>placebo).** Voxelwise correlations (TFCE, $p<0.0125$ FWE) between visual analogue “anxious” (A&B), “paranoid” (C&D) and “relaxed” (E&F) scores and CB1 and CB2 enriched connectivity. Red denotes positive, blue negative associations. Scatter plots show average correlations in clusters identified as significant and are for visualisation purposes only.

**Supplementary Table 2:** Regions showing significant voxelwise correlations between plasma cannabinoid levels and receptor-enriched functional connectivity in the THC + CBD and THC-only conditions.

| Receptor | Drug condition | Variable | Negative Correlations | Positive Correlations |
|---|---|---|---|---|
| CB1 (PET) | CBD | CBD levels | Frontal Orbital Cortex<br>Paracingulate Gyrus | Angular Gyrus<br>Central Opercular Cortex<br>Cingulate Gyrus<br>Cuneal Cortex<br>Frontal Opercular Cortex<br>Frontal Orbital Cortex<br>Frontal Pole<br>Heschl's Gyrus<br>Inferior Frontal Gyrus |

| | | | | |
|---|---|---|---|---|
| | | | | Insular Cortex<br>Juxtapositional Lobule Cortex<br>Lateral Occipital Cortex<br>Left Putamen<br>Middle Frontal Gyrus<br>Middle Temporal Gyrus<br>Parietal Opercular Cortex<br>Planum Polare<br>Planum Temporale<br>Postcentral Gyrus<br>Precentral Gyrus<br>Precuneous Cortex<br>Right Amygdala<br>Superior Frontal Gyrus<br>Superior Parietal Lobule<br>Superior Temporal Gyrus<br>Supramarginal Gyrus<br>Temporal Pole |
| CB1 (PET) | CBD | THC levels | Paracingulate Gyrus | Angular Gyrus<br>Central Opercular Cortex<br>Cingulate Gyrus<br>Frontal Opercular Cortex<br>Frontal Orbital Cortex<br>Frontal Pole<br>Heschl's Gyrus<br>Inferior Frontal Gyrus<br>Insular Cortex<br>Juxtapositional Lobule Cortex<br>Left Putamen<br>Middle Frontal Gyrus<br>Middle Temporal Gyrus<br>Paracingulate Gyrus<br>Parietal Opercular Cortex<br>Planum Polare<br>Planum Temporale<br>Postcentral Gyrus<br>Precentral Gyrus<br>Precuneous Cortex<br>Superior Frontal Gyrus<br>Superior Parietal Lobule<br>Superior Temporal Gyrus<br>Supramarginal Gyrus<br>Temporal Pole |
| CB1 (PET) | THC | THC levels | Middle Frontal Gyrus<br>Planum Temporale | Frontal Opercular Cortex<br>Heschl's Gyrus |

| | | | | |
|---|---|---|---|---|
| | | | Superior Temporal Gyrus<br>Supramarginal Gyrus | Insular Cortex<br>Parietal Opercular Cortex<br>Planum Temporale |
| CB2 (gene) | CBD | CBD levels | Frontal Orbital Cortex<br>Frontal Pole<br><br>Middle Frontal Gyrus<br>Parahippocampal Gyrus Temporal Pole | Central Opercular Cortex<br>Cingulate Gyrus<br>Frontal Opercular Cortex<br>Frontal Orbital Cortex<br>Frontal Pole<br>Heschl's Gyrus<br>Inferior Frontal Gyrus<br>Insular Cortex<br>Juxtapositional Lobule Cortex<br>Middle Frontal Gyrus<br>Middle Temporal Gyrus<br>Parietal Opercular Cortex<br>Planum Polare<br>Planum Temporale<br>Postcentral Gyrus<br>Precentral Gyrus<br>Superior Frontal Gyrus<br>Superior Parietal Lobule<br>Superior Temporal Gyrus<br>Supramarginal Gyrus<br>Temporal Pole |
| CB2 (gene) | CBD | THC levels | Frontal Orbital Cortex Frontal Pole<br>Middle Frontal Gyrus | Central Opercular Cortex<br>Cingulate Gyrus<br>Frontal Opercular Cortex<br>Frontal Orbital Cortex<br>Frontal Pole<br>Heschl's Gyrus<br>Inferior Frontal Gyrus<br>Insular Cortex<br>Juxtapositional Lobule Cortex<br>Middle Frontal Gyrus<br>Middle Temporal Gyrus<br>Paracingulate Gyrus<br>Parietal Opercular Cortex<br>Planum Polare<br>Planum Temporale<br>Postcentral Gyrus<br>Precentral Gyrus<br>Superior Frontal Gyrus<br>Superior Parietal Lobule<br>Superior Temporal Gyrus<br>Supramarginal Gyrus |

| | | | | |
|---|---|---|---|---|
| | | | | Temporal Fusiform Cortex<br>Temporal Pole |
| CB2 (gene) | THC | THC levels | | Central Opercular Cortex<br>Inferior Frontal Gyrus<br>Insular Cortex<br>Postcentral Gyrus<br>Precentral Gyrus |

**Supplementary Table 3:** Regions showing significant voxelwise correlations between subjective drug effect scores measured on a 0-10 visual analogue scale and receptor-enriched functional connectivity in the THC + CBD and THC-only conditions.

| Receptor | Drug condition | Subjective drug effect scale | negative | positive |
|---|---|---|---|---|
| CB1 (PET) | CBD | feel drug | Central Opercular Cortex<br>Cuneal Cortex<br>Frontal Opercular Cortex<br>Heschl's Gyrus<br>Insular Cortex<br>Juxtapositional Lobule Cortex<br>Parietal Opercular Cortex<br>Planum Temporale<br>Postcentral Gyrus<br>Supramarginal Gyrus | Cingulate Gyrus<br>Inferior Frontal Gyrus<br>Middle Frontal Gyrus |
| CB1 (PET) | THC | feel drug | | |
| CB2 (gene) | CBD | feel drug | Angular Gyrus<br>Cingulate Gyrus<br>Frontal Pole<br>Heschl's Gyrus<br>Middle Frontal Gyrus<br>Middle Temporal Gyrus<br>Paracingulate Gyrus<br>Planum Polare<br>Planum Temporale<br>Postcentral Gyrus<br>Precentral Gyrus<br>Superior Frontal Gyrus<br>Superior Temporal Gyrus<br>Supramarginal Gyrus<br>Temporal Pole | Postcentral Gyrus<br>Precentral Gyrus |
| CB2 (gene) | THC | feel drug | | |
| CB1 (PET) | CBD | anxious | Cingulate Gyrus<br>Frontal Orbital Cortex<br>Middle Frontal Gyrus | Central Opercular Cortex<br>Insular Cortex |

| | | | | |
|---|---|---|---|---|
| | | | Paracingulate Gyrus<br>Superior Frontal Gyrus | Juxtapositional Lobule Cortex<br>Middle Frontal Gyrus<br>Parietal Opercular Cortex<br>Planum Temporale<br>Postcentral Gyrus<br>Precentral Gyrus<br>Precuneous Cortex<br>Superior Parietal Lobule<br>Supramarginal Gyrus |
| CB1 (PET) | THC | anxious | Frontal Orbital Cortex<br>Precentral Gyrus | |
| CB2 (gene) | CBD | anxious | Cingulate Gyrus<br>Frontal Orbital Cortex<br>Frontal Pole<br>Middle Frontal Gyrus<br>Paracingulate Gyrus<br>Postcentral Gyrus<br>Precentral Gyrus<br>Superior Frontal Gyrus | |
| CB2 (gene) | THC | anxious | Juxtapositional Lobule Cortex<br>Postcentral Gyrus<br>Precentral Gyrus | |
| CB1 (PET) | CBD | paranoid | Central Opercular Cortex<br>Frontal Orbital Cortex<br>Frontal Pole<br>Heschl's Gyrus<br>Inferior Frontal Gyrus<br>Insular Cortex<br>Middle Frontal Gyrus<br>Middle Temporal Gyrus<br>Parietal Opercular Cortex<br>Planum Polare<br>Planum Temporale<br>Postcentral Gyrus<br>Precentral Gyrus<br>Superior Temporal Gyrus<br>Supramarginal Gyrus<br>Temporal Pole | |
| CB1 (PET) | THC | paranoid | Frontal Opercular Cortex<br>Frontal Orbital Cortex<br>Inferior Frontal Gyrus<br>Insular Cortex<br>Left Putamen<br>Middle Frontal Gyrus | |

| | | | | |
|---|---|---|---|---|
| CB2 (gene) | CBD | paranoid | Central Opercular Cortex<br>Frontal Opercular Cortex<br>Heschl's Gyrus<br>Inferior Frontal Gyrus<br>Insular Cortex<br>Planum Polare<br>Planum Temporale<br>Postcentral Gyrus<br>Precentral Gyrus<br>Supramarginal Gyrus<br>Temporal Pole | |
| CB2 (gene) | THC | paranoid | Postcentral Gyrus<br>Precentral Gyrus | |
| CB1 (PET) | CBD | relaxed | Central Opercular Cortex<br>Insular Cortex<br>Parietal Opercular Cortex<br>Planum Temporale<br>Postcentral Gyrus<br>Precuneous Cortex<br>Right Amygdala<br>Superior Parietal Lobule | Cingulate Gyrus<br>Frontal Pole<br>Paracingulate Gyrus<br>Superior Frontal Gyrus |
| CB1 (PET) | THC | relaxed | Central Opercular Cortex<br>Frontal Opercular Cortex<br>Frontal Orbital Cortex<br>Frontal Pole<br>Inferior Frontal Gyrus<br>Insular Cortex<br>Left Putamen<br>Paracingulate Gyrus<br>Precentral Gyrus<br>Superior Frontal Gyrus | |
| CB2 (gene) | CBD | relaxed | | Temporal Pole |
| CB2 (gene) | THC | relaxed | Planum Polare<br>Postcentral Gyrus<br>Precentral Gyrus<br>Superior Frontal Gyrus<br>Superior Parietal Lobule | |
